# Effects of H2A.B incorporation on nucleosome structures and dynamics

**DOI:** 10.1101/2020.06.25.172130

**Authors:** Havva Kohestani, Jeff Wereszczynski

## Abstract

The H2A.B histone variant is an epigenetic regulator involved in transcriptional upregulation, DNA synthesis, and splicing that functions by replacing the canonical H2A histone in the nucleosome core particle. Introduction of H2A.B results in less compact nucleosome states with increased DNA unwinding and accessibility at the nucleosomal entry and exit sites. Despite being well characterized experimentally, the molecular mechanisms by which H2A.B incorporation alters nucleosome stability and dynamics remain poorly understood. To study the molecular mechanisms of H2A.B, we have performed a series of conventional and enhanced sampling molecular dynamics simulation of H2A.B and canonical H2A containing nucleosomes. Results of conventional simulations show that H2A.B weakens protein/protein and protein/DNA interactions at specific locations throughout the nucleosome. These weakened interactions result in significantly more DNA opening from both the entry and exit sites in enhanced sampling simulations. Furthermore, free energy profiles show that H2A.B containing nucleosomes have significantly broader free wells, and that H2A.B allows for sampling of states with increased DNA breathing, which are shown to be stable on the hundreds of nanoseconds timescale with further conventional simulations. Together, our results show the molecular mechanisms by which H2A.B creates less compacted nucleosome states as a means of increasing genetic accessibility and gene transcription.

**SIGNIFICANCE:** Nature has evolved a plethora of mechanisms for altering the physical and chemical properties of chromatin fibers as a means of controlling gene expression. These epigenetic processes may serve to increase or decrease DNA accessibility, manage the recruitment of remodeling factors, or tune the stability of the nucleosomes that make up chromatin. Here, we have used both conventional and enhanced sampling molecular dynamics simulations to understand how one of these epigenetic mechanisms, the substitution of canonical H2A proteins with the H2A.B variant, exerts its influence on the structures and dynamics of the nucleosome. Results show at the molecular level how this variant alters inter-molecular interactions to increase DNA accessibility as a means of increasing genetic accessibility and gene transcription.

## INTRODUCTION

The capability to store, transfer, and modify genetic information is one of the distinctive features of living organisms. Eukaryotes have solved this problem by evolving multiple molecular mechanisms to control the packing and accessibility of their genetic code in their confined cellular environments. Central to these processes is the nucleosome core particle (NCP), a complex composed of eight histone proteins that wraps ∼147 DNA basepairs (1–4). It consists of eight proteins, including two copies each of the H2A, H2B, H3, and H4 histones. Each core histone is comprised of three α-helices (α_1_, α_2_, and α_3_) connected by two short loops (L1 and L2), with each histone also having various lengths of unstructured N- and C-terminal “tail” regions. The histones are arranged into a (H3-H4)_2_ tetramer that is flanked by two H2A-H2B dimers, (Fig. 1) (1–7). Nucleosomes assemble on long DNA molecules to form chromatin fibers which condense into higher-order structures that are stabilized by significant electrostatic forces between the DNA and histone protein core (4, 8). Several mechanisms exist to tune the nucleosome stability, and thus DNA accessibility and chromatin structure, which include post-translational modifications to select histone residues, the recruitment of ATP-dependent chromatin remodeling complexes, and the substitution of canonical histones with histone variant proteins (9–19).

**Figure 1:**
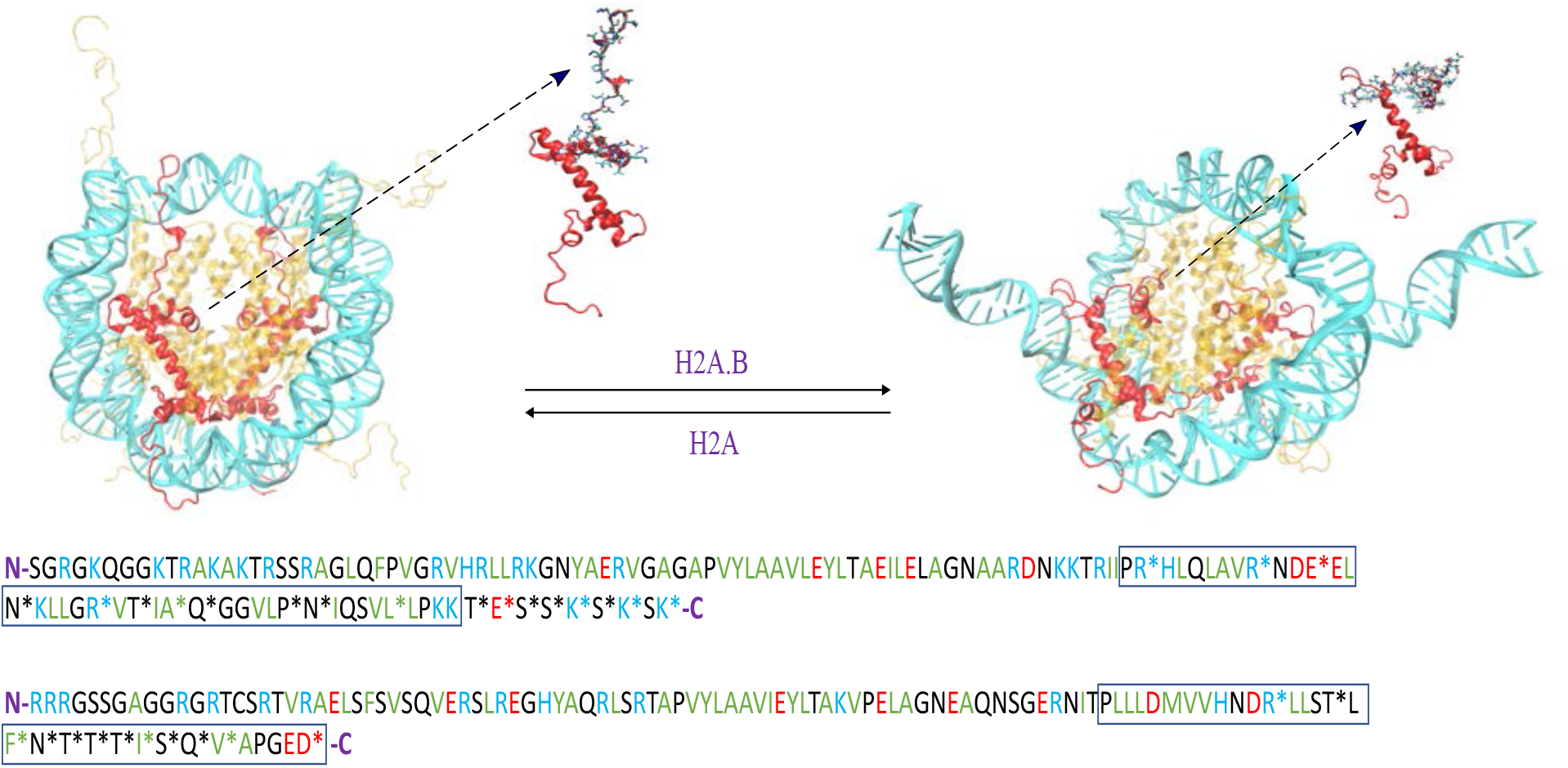
Substitution of canonical H2A with H2A.B results in more open nucleosome structures (top). Sequence alignment of H2A and H2A.B histones (bottom) show H2A.B has a shorter docking domain (boxed), alterations to the acidic patch, and no C-terminal tails (11). Basic residues are in blue, acidic in red, hydrophobic in green, polar in black.

The H2A histone family has the most diverse and populated group of variants (15, 20–23). This group includes the canonical H2A histone as well as the H2A.B, H2A.L, H2A.P, H2A.W, H2A.Z, H2A.X, and macroH2A variants(24). Of particular note, H2A.B (which was formerly referred to as H2A.Bbd for Barr-body-deficient) is associated with DNA synthesis, splicing, and transcription upregulation (23–29). It was first described by Chadwick and Willard as a histone variant that is excluded from the inactive X chromosome of females and is localized to H4 hyperacetylated regions (30). It is only 48-50% identical to canonical H2A, lacks both the C-terminal tail and parts of the “docking domain,” and does not contain the “acidic patch” which promotes chromatin condensation (31–33).

Substitution of canonical H2A with H2A.B results in large-scale structural changes within the nucleosome. *In vitro*, H2A.B containing nucleosomes protect only 118-130 basepairs from micrococcal nuclease digestion, suggesting that the entry and exit DNA segments are more exposed to solution in H2A.B nucleosomes (34, 35). Fluorescence resonance energy transfer (FRET) experiments have also shown this, with H2A.B substitution increasing the distance between DNA ends in reconstituted nucleosomes (34). Both atomic force and cryo-electron microscopy experiments have revealed coarse representations of these extended states, with the most common conformations observed to have ∼10 basepairs of DNA unwrapped from each DNA end (35). These extended states were further confirmed with small angle X-ray scattering experiments, which showed that in solution H2A.B substitution increases the nucleosome’s radius of gyration from 43.4±0.3 Å to 49.4±0.4 Å (29).

Together, these results demonstrate that H2A.B has a significant affect on nucleosome structures which directly influences their functions. Indeed, the most comprehensive experimental atomic-scale study of H2A.B’s effect is a recently published crystal structure of the H2A.B-H2B dimer at 2.6 Å resolution which demonstrates that variant substitution drastically alters the surface electrostatic potential of the dimer by eliminating the overall positive charge in the DNA binding site (36). However, there has yet to be a structure solved of a complete H2A.B containing nucleosome, likely due to the absence of the C-terminal, shorter docking domain, and increased DNA flexibility hindering crystallization. This has precluded further studies on the effects of H2A.B substitution on nucleosome dynamics, and how those effects propagate into large-scale structural rearrangements and increased DNA motions. To address these challenges, we have constructed an atomic-scale model of an H2A.B containing nucleosome. Using comparative conventional and free energy molecular dynamics simulations, our results demonstrate that H2A.B substitution alters interactions at the DNA-dimer and dimer-tetramer interfaces. These changes show distinct energetic characteristics, with modifications in the L2 loops leading to increased DNA opening and changes in the docking domains likely resulting in destabilization of the dimer/tetramer interface on timescales significantly longer than those sampled here. Taken together, our results show how histone substitution creates less compacted nucleosomes which are responsible for H2A.B’s biological functions.

## MATERIALS AND METHODS

### Molecular Dynamics Simulations

A model of the canonical nucleosome was constructed based on the 1KX5 crystal structure (3). The H2A.B containing nucleosome was modelled by modifying the H2A sequences in the 1KX5 structure to the H2A.B sequence (11). After initiation of this work, a structure of the H2A.B/H2B dimer was released, which showed a good agreement to our model, with a heavy atom root-mean square deviation (RMSD) of 1.8 Å between the two structures, indicating our model was of sufficient accuracy for molecular dynamics simulations (36). All systems were built in the tleap tool of the Amber molecular dynamics package (37). The Amber ff14SB, OL15, and TIP3P force fields were used for the protein, DNA, and water force fields, while ions parameters were based on the Joung and Cheatham model (38–41). Systems were neutralized and solvated in a water box with a 12 Å buffer of solvent and an additional a 0.15 M NaCl concentration. All systems were minimized twice for 10000 steps, once with and once without heavy atom restraints. Systems were gradually heated in the canonical ensemble, from 0 K to 300 K over 5000 steps with restraints on solute heavy atoms. The restraints were released by relaxing the systems in the isothermal-isobaric ensemble. The Langevin piston was used for a barostat and Langevin dynamics were performed for temperature control with a collision frequency of 2 ps^−1^ (42). Systems of the compact and open states were each performed twice for 550 ns, resulting in 8.8 µs of explicit solvent simulations. All simulations was performed by with the GPU accelerated version of PMEMD (43).

Enhanced sampling simulations followed a similar protocol with three major exceptions. First, the H3 N-terminal tails were truncated. Second, they were performed in implicit solvent while the default PBRadii of the Generalized Born Surface Area model was set to *mbondi2* with *igb=5* and a salt concentration of 0.15 M (44, 45). This implicit solvent model has previously shown good agreement with experimental results for nucleosome systems, including for million atom-scale chromatin fiber models which are characterized by linker DNA and histone tail motions, as well as sub-nucleosomal particles that have extensive DNA opening motions (46). We used the default value of rgbmax=25 for the maximum distance between atom pairs to calculate the Born radii and a nonbonded cutoff of effectively infinity (999.0 Å). The ParmEd (parameter file editor) program was used to repartition the hydrogen atom masses to reduce the system high frequency motions, which allowed the use of a 4 ps MD timestep (47). Finally, Adaptively Biased Molecular Dynamics (ABMD) were performed (48). ABMD belongs to a class of enhanced sampling methods, which include methods such as metadynamics and adaptive biasing force, in which an additional potential is added along a low number of reaction coordinate to enhance sampling (49, 50). This potential is modified as the simulation progresses to adaptively flatten the underlying free energy surface, and in ABMD this potential is integrated to compute the potential of mean force along the desired reaction coordinate(s). Here, we have implemented ABMD along two reactions coordinates: the first being the radius of gyration of the heavy atoms of the first 73 DNA basepairs, and the second as the radius of gyration of the heavy atoms of the last 73 DNA basepairs. This choice was motivated by initial tests which showed that choosing only the radius of gyration of the entire DNA molecule resulted in highly degenerate structures. A flooding time of 100 ps was used along with a spatial resolution of 1 Å. For each system, three simulations of 1.6 µs were run, resulting in 4.8 µs of sampling for each. Simulations were initiated from the closed DNA states, with an equilibration time of 50 ns. A summary of the simulations performed here is included in Table S1.

### Simulation Analyses

We used the cpptraj clustering tool to extract the dominant structural clusters. The optimal number of clusters was based on Davis-Bouldin Index (DBI) and seudo-F statistic (pSF) metrics (51, 52). The maximum DBI and minimum pSF numbers gives the optimal number of structural clusters. This number is six for both H2A and H2A.B NCPs (Fig. S1). UCSF Chimera modeller was used to reattach the truncated H3 histone tails for the final explicit solvent simulations of open states (53, 54).

To estimate interaction energies, we utilized an MM/GBSA approach.(55, 56). In this method, the change in energy between two states is estimated through an endpoint approach:

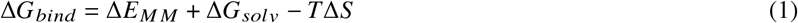

Where the overall free energy of binding (ΔG) is computed from the differences in gas-phase molecular mechanics energies (Δ*E*_*MM*_), solvation free energy differences (Δ*G*_*solv*_), and conformational entropy (Δ*S*) between states. Here, we have employed the “one-trajectory” approach in which differences between the “bound” and “unbound” states are extracted from the same simulations, thus the internal energy components of the Δ*E* term are zero and the only contributing terms are from the van der Waals and electrostatics. The solvation term is composed of two portions: a polar and an apolar term. For the polar term we used the Generalized Born model with the *igb*=8 and *igb*=5 implicit solvent models in a salt concentration of 0.15 M with the *mbondi3* and *mbondi2* radii set respectively (45, 57). Apolar solvation energies were calculated based on the change of solvent exposed surface area (SASA) between states scaled by a surface tension of 0.0072 kcal/mol/Å^2^. Comparison of the apolar SASA based methods used here with an alternative method based on a cavity and dispersion term show little difference between the two methods (Table S2), indicating that the overall conclusions are robust to the SASA methods used. The change in conformational entropy was excluded due to convergence issues.

Although MM/GBSA values contain an estimate of solvation entropic differences, they do not contain multiple important entropic terms such as the conformational entropy of the solute or explicit solvent molecules. Therefore, they are closer to an enthalpic difference then a free energy difference, and we have labeled them as ΔE differences, although we emphasize that they do not represent a purely enthalpic difference between states. In addition, for clarity we have grouped the energy differences into two terms: Δ*E*_*vdW*_ and Δ*E*_*elec*_. Δ*E*_*vdW*_ is the sum of the molecular mechanics van der Waals energy and the solvation apolar energy differences, and *E*_*elec*_ is the sum of the molecular mechanics electrostatics and polar solvation energy differences.

The *MMPBSA*.*py* script was used to perform an MM/GBSA analysis (54). This was performed on the final 500 ns of the compact state simulations. This analysis was also performed for the time intervals of 250 ns and 500 ns to evaluate the conformational convergence. Binding energies were calculated and averaged based on 100 frames from each of the two trajectories for a system, with each frame having two copies of the DNA/dimer, dimers-tetramer, and H2A/H2A.B-tetramer interfaces, resulting in 400 frames for each calculation. The errors are reported as standard error of the mean with a 5 ns decorrelation time. The cpptraj analysis tool was used for root-mean-square-deviation (RMSD), Radius of gyration (Rg), and structural clustering analyses (58). Visual Molecular Dynamics (VMD) was used for figures (59). Correlation analyses were performed with the *g-Correlation* program on the and 1 atoms for every five ps of the input trajectories (60).

## RESULTS

### Simulations of Compact States Show Weaker Interfaces in H2A.**B Nucleosomes**

Two, 550 ns conventional simulations of canonical and H2A.B containing nucleosomes were performed based upon the compact state in the 1KX5 crystal structure. Large scale dynamics were similar between systems, which were primarily characterized by H3 tail collapse onto the nucleosomal DNA, as had been observed in other full-size nucleosome simulations (61–66). This is exemplified by root-mean-square-deviation (RMSD) calculations, which show changes on the order of 1.0 and 0.5 Å in the DNA and core protein structures, respectively, in these systems (Figs. S2 & S3). Given that experiments have suggested that open states of H2A.B nucleosomes dominate in solution, the fact that these simulations remained in compact states is likely an artifact of the limited sampling inherent to conventional MD simulations.

Although large-scale structural rearrangements were not observed in these simulations, energetic analysis shows that incorporation of H2A.B may lead to destabilization of the compact nucleosome structure. An MM/GBSA analysis shows that dimer-tetramer interactions in canonical nucleosomes are largely stabilized by favorable van der Waals (vdW) interactions, which are counterbalanced by non-favorable electrostatic interactions between the two cationic species (Table 1). In H2A.B containing nucleosomes, there are reduced dimer-tetramer contacts, which results in a weaker interface. Although the electrostatic repulsive energies are reduced, the attractive vdW forces are much more significantly reduced, resulting in a less favorable Δ*E*_*total*_. In both systems we note that nucleosome stability is enhanced by strong DNA-dimer interactions, which are roughly equal in strength in H2A and H2A.B containing systems. This is in line with experiments which have shown that DNA is required for nucleosome stability at physiological salt concentrations, and that the histone core will only assemble at high salt concentrations without DNA (67–69). We note that the MM/GBSA analysis presented here is relatively crude and does not include important thermodynamic terms such as conformational entropy and explicit solvation, therefore although the trends are of qualitative value, care should be taken in quantitative interpretation of the energetic values (56, 70–73). Interaction energies were similar regardless of the generalized Born parameters that were used (Table S3), whether only the first half or the complete trajectories were used (Table S4), or whether the nonpolar term was computed based on only the change in solvent accessible surface area or if a cavity-dispersion term was used (Table S2) (74, 75).

**Table 1:**
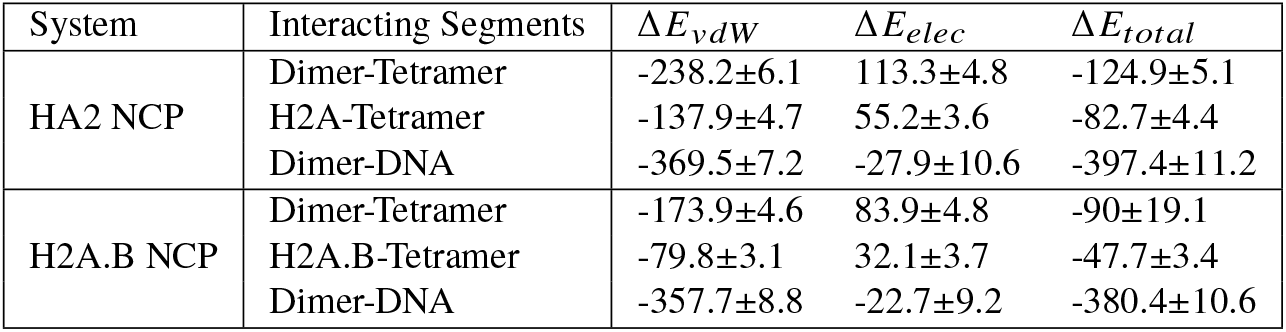
Energy differences (kcal/mol) of association between the dimer/tetramer, H2A/tetramer, and dimer/DNA constituents as estimated by an MM/GBSA analysis. A negative value indicates more favorable interactions in the complex relative to in solution. The total interaction energy (Δ*E*_*t tal*_) is a sum of the van der Waals (Δ*E*_*vdW*_) and electrostatic (Δ*E*_*elec*_) interactions.

To determine the location of residues that contribute to destabilization of H2A.B nucleosomes, an energetic decomposition was performed for residues in the histone core. Residues in the long histone tails were excluded from this analyses, as their sampling is highly stochastic and subject to significant noise, which may create unconverged results at the single residue level (76). The change in energy upon H2A.B substitution, ΔΔ*E*_*total*_ = Δ*E*_*T otal,H*2*A*.*B*_ −Δ*E*_*T otal,H*2*A*_, was computed, and the most significant values are reported in Table 2. Residues that contribute the most to canonical dimer-tetramer stability are primarily located in three regions: the H2B α_2_/H4, the H2A/H3-H4, and the H2A docking domain/H3 interfaces, which we denote as regions one, two, and three in Fig. 2 (Tables 2, S5 & S6). Upon H2A.B substitution, these interaction energies are altered (Tables S7 & S8), and residues in these three areas primarily lead to destabilization of the histone core (Table 2). The first region of this interface corresponds largely to residues Asp68, Asp85, Lys91, Arg92, and Tyr98 from the H4 α_3_ helix and Glu73 and Leu97 from the H2B α_2_ helix. Together, substitution destabilizes this complex through a ΔΔ*E*_1_ = 6.6 ± 2.6 kcal/mol of energy. The second region corresponds primarily to residues Glu94 and Glu97 of the H3 α_2_ helix, and Arg134 of the H3 C-terminal, and Thr101 and Ala103 of the H2A docking domain. It has an energy difference of ΔΔ*E*_2_ = 1.1 ± 1.2 kcal/mol, which is not statistically significant but does suggest there may be a relatively minor destabilization in this region. The most significant energy difference is in region three, which includes Arg99, Leu115, Lys118, and Lys119 of the H2A docking domain, Glu121 of the H2A C-terminal, Asn108 of the H3 α_2_ helix, and Lys44 of the H4 L1-loop. Residues Leu115 through Glu121 are absent in H2A.B while Arg99 of H2A is mutated to a threonine. This results in a significant destabilization of the dimer-tetramer interface in this region, with an energy difference of ΔΔ*E*_3_ = 13.3 ± 3.8 kcal/mol.

**Table 2:**
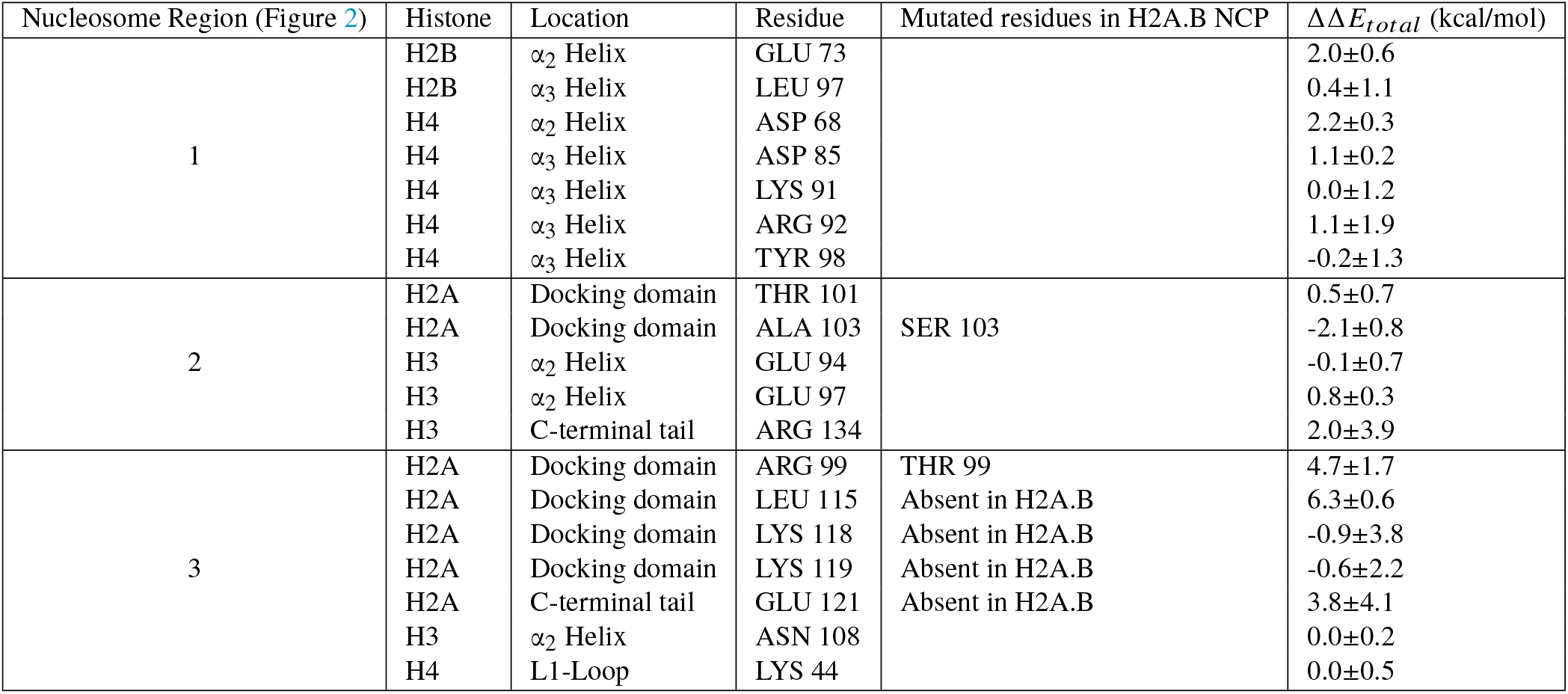
Residues with the most significant differences in interaction energies in the three regions identified in Figure 2 at the dimer/tetramer interface (Δ Δ*E*) between H2A and H2A.B containing nucleosomes. Negative values correspond to locations that are more favorable in H2A.B systems, whereas positive values are more favorable in H2A nucleosomes.

**Figure 2:**
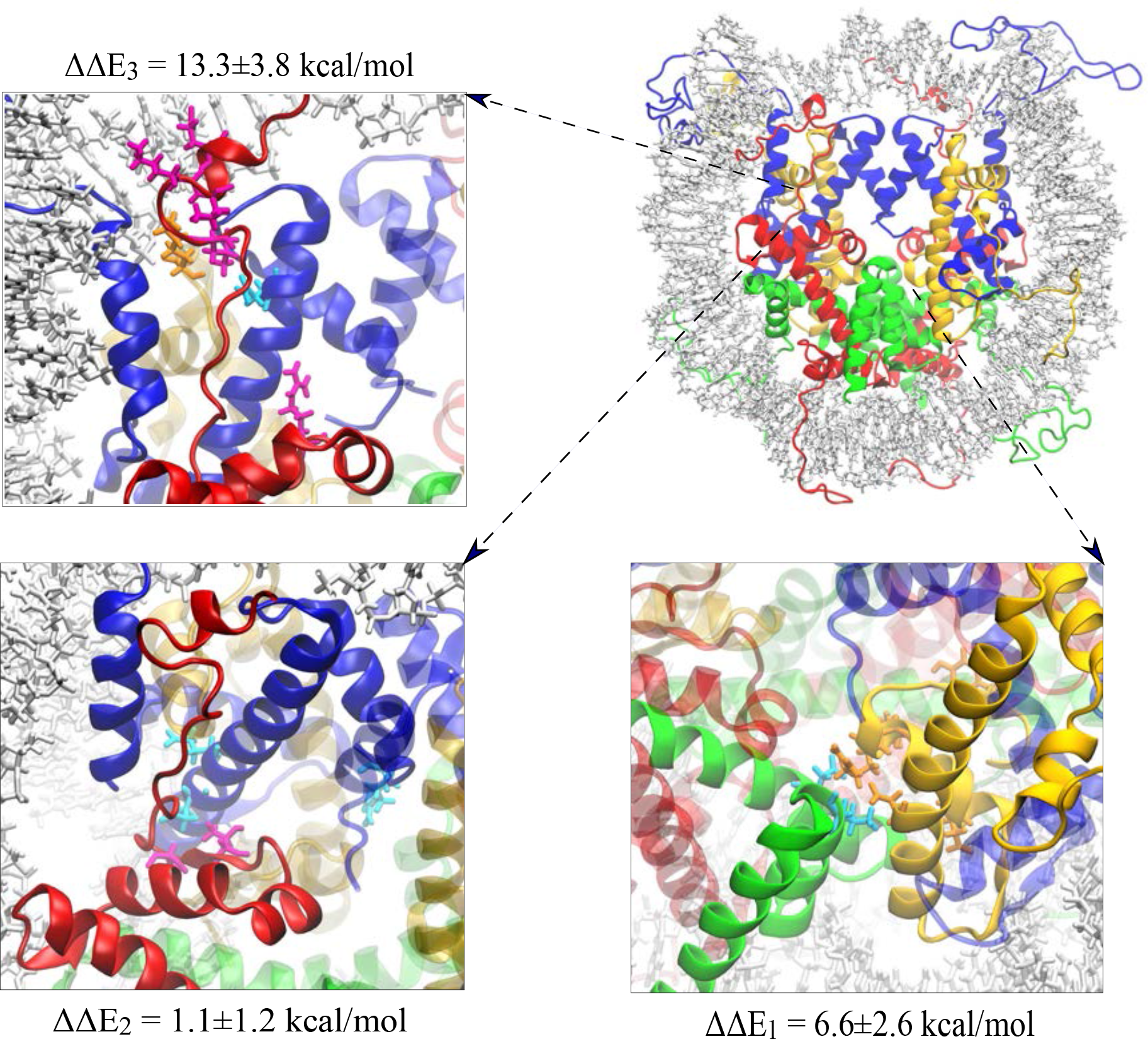
Nucleosome regions that have the most significant change in interaction energies upon H2A.B incorporation at the dimer/tetramer interface, along with their energy changes (kcal/mol). These regions correspond to (1) the H2B α_2_/H4, (2) the H2A/H3-H4, and (3) the H2A docking domain/H3 interfaces.

A similar energetic decomposition was performed for residues at the dimer core/DNA interface. In canonical nucleosomes, the major points of contacts between H2A and DNA occur in the α_1_ helix, the L1, and the L2 loops, along with Arg17 in the αN helix near the N-terminal tail (Fig. 3). In each of these regions there are strong stabilizing contacts formed, particularly between Arg42, Thr76, and Arg77 with the DNA (Table 3). Substitution of H2A.B abolishes both the α_1_ and L2 loop contacts while maintaining the L1 contacts (Table 4). This results in roughly similar interactions at the base of the nucleosome opposite of the dyad, but reduced contacts closer to the entry and exit DNA sites (Fig. 3). Notably, by changing the L2 loops from having a net charge of +2 to neutral upon H2A.B substitution, the DNA interaction energy of these loops is significantly destabilized from −10.7±1.5 kcal/mol to −3.1±1.9 kcal/mol. Contacts between H2B and DNA were similar in both systems, with stabilizing contacts largely formed between the L2 loop and DNA.

**Figure 3:**
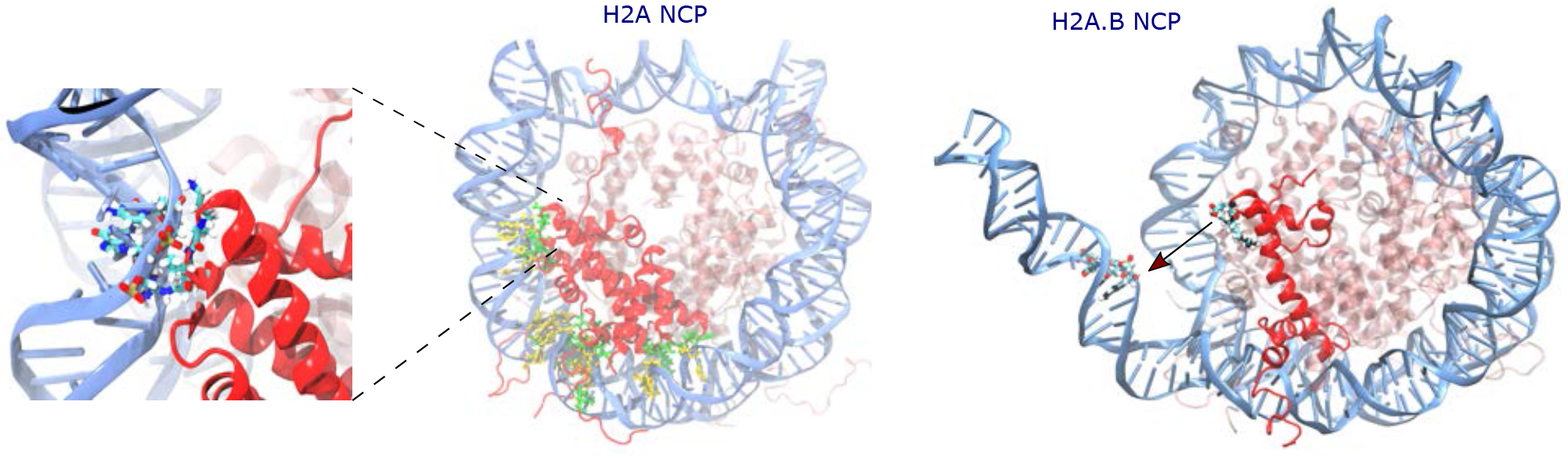
Stabilizing DNA/dimer contacts in H2A and H2A.B containing systems. Protein residues and DNA basepairs forming contacts are shown in green and yellow, respectively. Residues at the H2A/H2A.B L2 loops (see zoomed-in figure on the left) have the largest difference in stabilizing contacts upon variant substitution. DNA is detached from protein core in H2A.B NCP as a result of weakened DNA-L2 loop bonds. This is in agreement with the increased distance at this interface in H2A.B NCP, see Fig. 6.

**Table 3:** Residues with the most significant interaction energies at the dimer/DNA interface in H2A systems (kcal/mol). Negative values correspond to interactions that stabilize DNA binding.

| H2A NCP |  |  |  |  |  |
| --- | --- | --- | --- | --- | --- |
| H2A Histone |  |  | H2B Histone |  |  |
| Residue | Location | $\Delta E_{total}$ (kcal/mol) | Residue | Location | $\Delta E_{total}$ (kcal/mol) |
| ARG 17 | $\alpha N$ Helix | $-7.4 \pm 1.1$ | SER 52 | L1 Loop | $-1.7 \pm 0.6$ |
| GLY 28 | $\alpha_1$ Helix | $-0.6 \pm 0.5$ | SER 53 | $\alpha_2$ Helix | $-3.2 \pm 1.1$ |
| ARG 29 | $\alpha_1$ Helix | $-3.1 \pm 1.2$ | ARG 83 | L2 Loop | $-4.7 \pm 1.2$ |
| ARG 42 | L1 Loop | $-4.9 \pm 1.9$ | SER 84 | L2 Loop | $-3.4 \pm 1.3$ |
| VAL 43 | L1 Loop | $-0.6 \pm 0.6$ | THR 85 | L2 loop | $-6.3 \pm 1.7$ |
| GLY 44 | L1 Loop | $-1 \pm 0.3$ | | | |
| LYS 75 | L2 Loop | $-1.8 \pm 1.4$ | | | |
| THR 76 | L2 Loop | $-4.5 \pm 1.5$ | | | |
| ARG 77 | L2 Loop | $-4.4 \pm 1.6$ | | | |

**Table 4:** Residues with the most significant interaction energies at the dimer/DNA interface in H2A.B systems (kcal/mol). Negative values correspond to interactions that stabilize DNA binding.

| H2A.B NCP |  |  |  |  |  |
| --- | --- | --- | --- | --- | --- |
| H2A.B Histone |  |  | H2B Histone |  |  |
| Residue | Location | $\Delta E_{total}$ (kcal/mol) | Residue | Location | $\Delta E_{total}$ (kcal/mol) |
| ARG 17 | αN Helix | $-7.7 \pm 0.9$ | SER 52 | L1 Loop | $-1.3 \pm 0.8$ |
| THR 18 | αN Helix | $-0.2 \pm 0.2$ | SER 53 | $\alpha_2$ Helix | $-2.9 \pm 1.2$ |
| ARG 42 | L1 Loop | $-4.7 \pm 1.2$ | ARG 83 | L2 Loop | $-4.4 \pm 1.1$ |
| LEU 43 | L1 Loop | $0.1 \pm 0.5$ | SER 84 | L2 Loop | $-3.7 \pm 1.6$ |
| SER 44 | L1 Loop | $-3 \pm 0.8$ | THR 85 | L2 loop | $-6 \pm 1.1$ |

### Enhanced Sampling Simulations Show Increased DNA Opening in H2A.**B Nucleosomes**

Adaptively biased MD (ABMD) simulations were performed to overcome the limited timescales of conventional MD simulations which precluded sampling of open nucleosomal states. For each system, three ABMD simulations of 1.6 µs each were performed. The reaction coordinates for these calculations were the radius of gyration (Rg) of the entry and exit DNA (see methods for details). To obtain sufficient sampling, simulations were performed in an implicit solvent environment, and the H3 N-terminal tails were truncated, as they tend to collapse on the DNA and prevent DNA breathing (76). Based on previously published and our own preliminary simulations, removal of the H3 N-terminal tails was necessary for allowing DNA opening on the microsecond timescale (77). However, the presence of the other histone tails did not significantly impair these motions, and they were therefore maintained in enhanced sampling simulations. Convergence of these simulations was assessed by Kullback–Leibler divergence calculations to the final probability states which demonstrated that in most simulations there was little change over the last 600 ns (Fig. S4) (78, 79). In the case of canonical nucleosomes, the maximum observed Rgs were 55 and 50 Å for the entry and exit DNA, indicating a moderate amount of DNA breathing (Fig. 4). The potential of mean force (PMF) shows a single free-energy well located at 44.5 and 41.3 Å (Fig. 5). This corresponds to states with slightly opened DNA structures, which we attribute to the implicit solvent environment and the lack of H3 tails. We note that in some of the one-dimensional projections of the ABMD sampling in Figure 4 there are regions of low fluctuations which correspond to metastable bound entry or exit DNA states. Specifically, these occurred when an asparagine near the H2A L2 loop inserted into the DNA and reduced it fluctuations. However, the resulting PMFs show that these are unfavorable states with high free energies, and we emphasize that the ABMD force is applied in two dimensions, and although the projections may appear to have low fluctuations in one dimension the systems fluctuate in the two dimensional plane and sample multiple ABMD grid points in these time periods.

**Figure 4:**
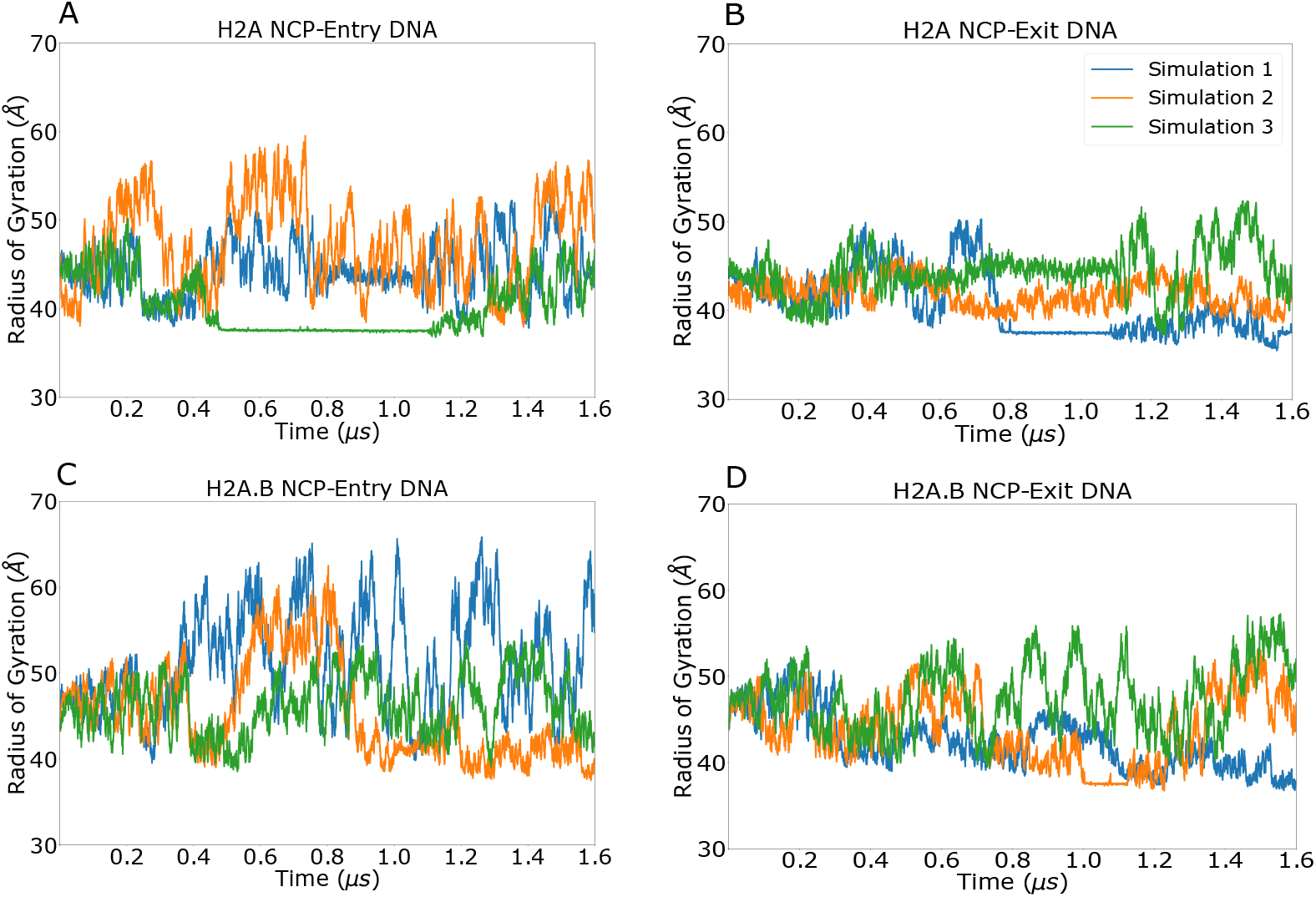
Radius of gyration for the entry and exit DNA segments in H2A and H2A.B nucleosomes during ABMD simulations. Both the entry and exit DNA segments in H2A.B systems have increased sampling and extensions relative to canonical nucleosomes which corresponds to more DNA breathing and increased DNA exposure. For representative structures at various radii of gyration, see Figs. 7 & 8.

**Figure 5:**
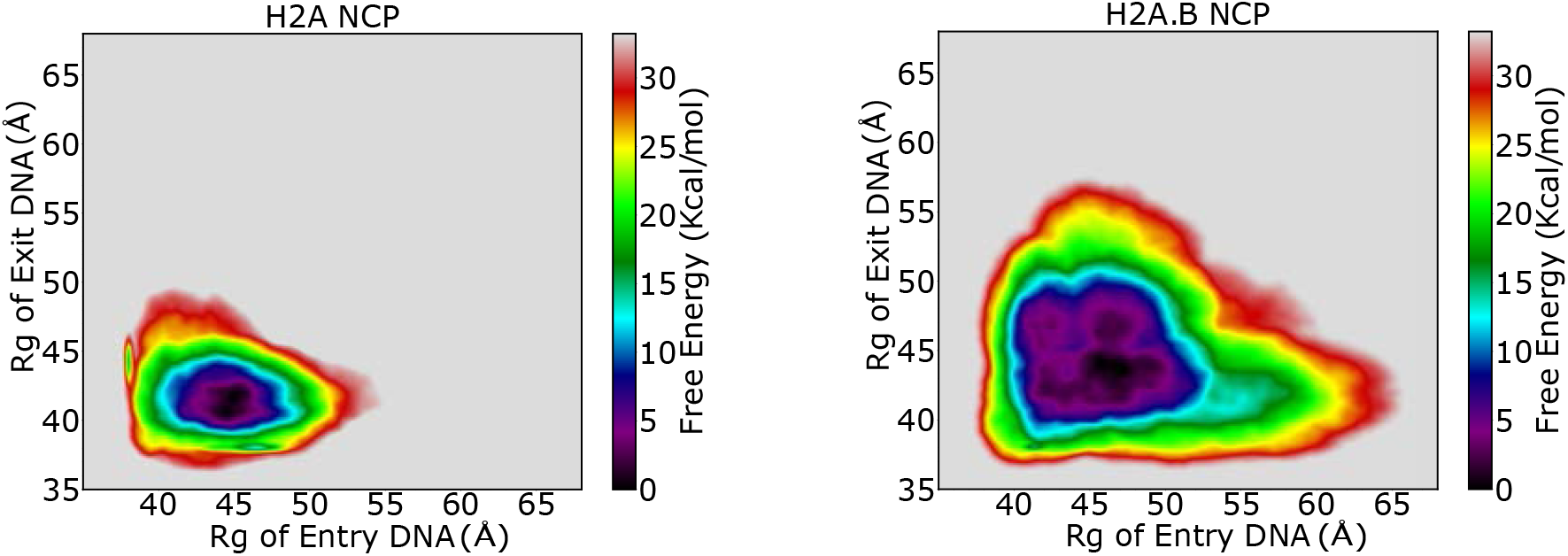
Free energy landscapes of canonical and H2A.B containing nucleosomes. Both systems have a single energy basin, however replacement of H2A with H2A.B results in a broader well and increased sampling of higher energy states.

In contrast, ABMD simulations with H2A.B nucleosomes experienced significantly more DNA opening. The maximum Rgs in these simulations were 66 and 57 Å for the entry and exit DNA, indicating that states with more DNA breathing from both the entry and exit sites were sampled (Fig. 4). Although the absolute location of the free energy minima in the PMFs is nearly identical to the canonical nucleosome (46.5 and 44.2 Å for entry and exit DNA) the free energy wells are significantly broader and extend to higher Rg values, with low free energy states extending from 40 to 50 Å in each dimension.

Energetic analyses of the closed states simulations indicated that DNA-dimer interactions were significantly reduced in the H2A.B L2 loops upon variant substitution. In ABMD simulations, this destabilization resulted in increased separation between the DNA and H2A.B L2 loops with DNA. Indeed, in each of the three canonical simulations, the distance between the DNA and L2 loops remained at its initial value of ∼13 Å, with the exception being the second ABMD simulation in which there were three detachment events where this distance increased (Fig. 6). In contrast, in each of the H2A.B ABMD simulations the DNA/L2 Loop distance experienced multiple opening events for both the entry and exit DNA segments.This effect directly resulted in the frequent opening of up to 20 DNA basepairs from each side of the nucleosome.

**Figure 6:**
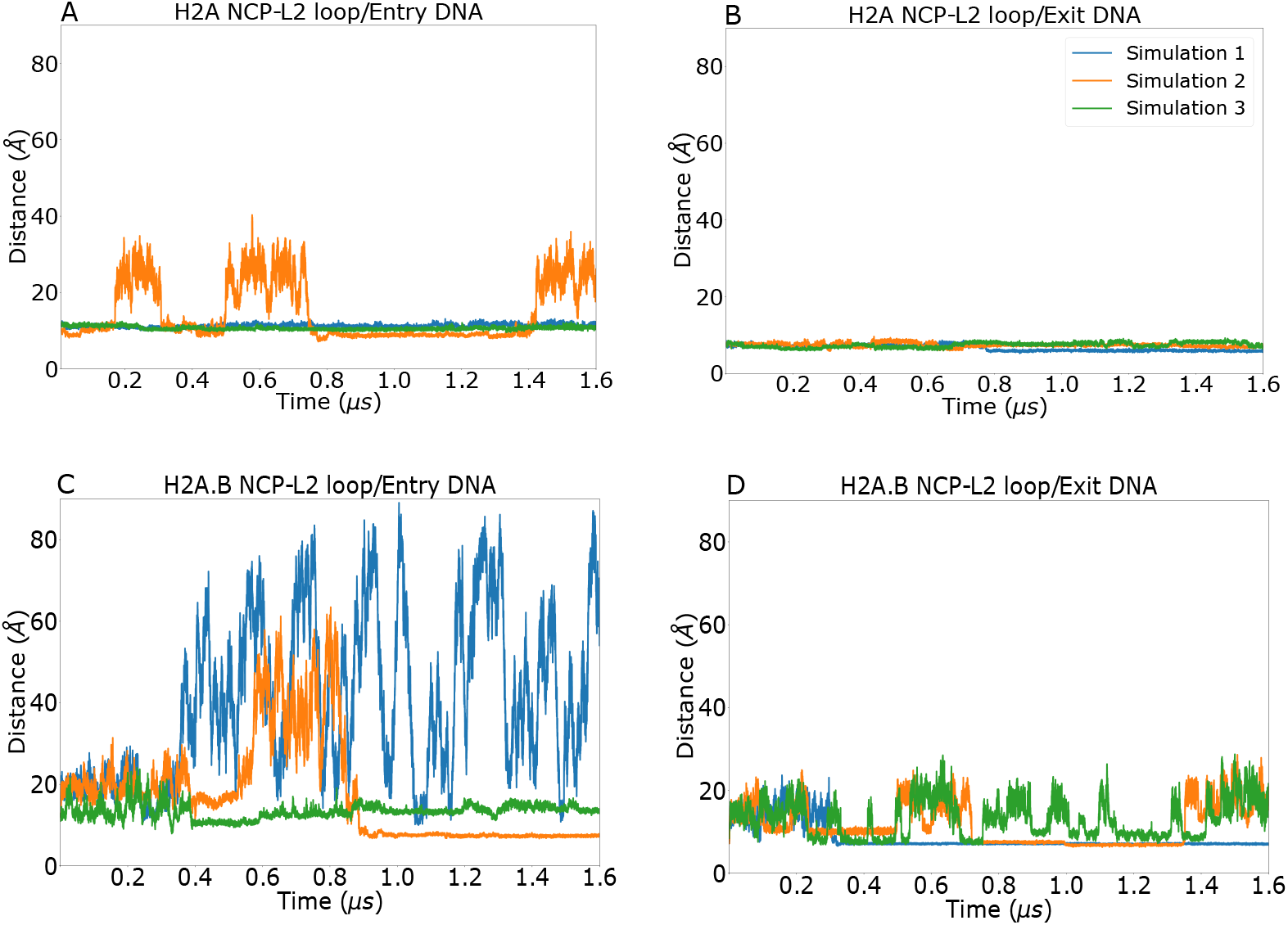
Separation distance between H2A (a&b) and H2A.B (c&d) L2 loops with their adjacent DNA basepairs in ABMD simulations. In canonical systems these distances are stable, indicating that the L2 loops remain in contact with the DNA throughout the majority of the simulations. In H2A.B systems there is a higher degree of detachment as the distances between the L2 loops and DNA sample a wide array of states throughout the simulations. For representative structures at various L2/DNA distances, see 7 & 8.

To visualize the sampled states, a clustering analysis was performed on the trajectories for each system. Both Davis-Bouldin Index (DBI) and pseudo-F statistics (pSF) indicate that six clusters are optimal for describing each system (Fig. S1). The representative structures from these clusters show that canonical nucleosomes adopt predominantly compact and semi-open states (Fig. 7). These states are located primarily in the PMF free energy well, and have DNA Rg values between 37.5 and 49.8 Å (Fig. S5). H2A.B containing nucleosome structures represent more open states, with Rg values ranging from 41.4 to 56.2 Å (Figs. 8 & S6). In both systems the histone core was relatively unchanged with RMSD values not exceeding 2 Å, demonstrating the histone core’s stability in these simulations (Fig. S11). ABMD simulations revealed the presence of open canonical and H2A.B nucleosome states. However, to achieve the required level of sampling, simulations had to be performed in implicit solvent and without the N-terminal H3 tails. To analyze the stability of these open states in more physiological conditions, the H3 tails were modelled onto the representative cluster centers from ABMD calculations and these structures were resolvated. For each structure, two 550 ns explicit MD simulations were performed resulting in 6.6 µs of sampling. In each simulation the histone core remained relatively stable, with RMSD values that did not exceed 4 Å (Fig. S12). In all open simulations the DNA remained in their initial open states as evidenced by stable radii of gyration for the entry and exit DNA segments (Fig. S13). Taken together, this confirms the stability of these open states in these more physiological conditions on the hundreds of nanoseconds timescale.

**Figure 7:**
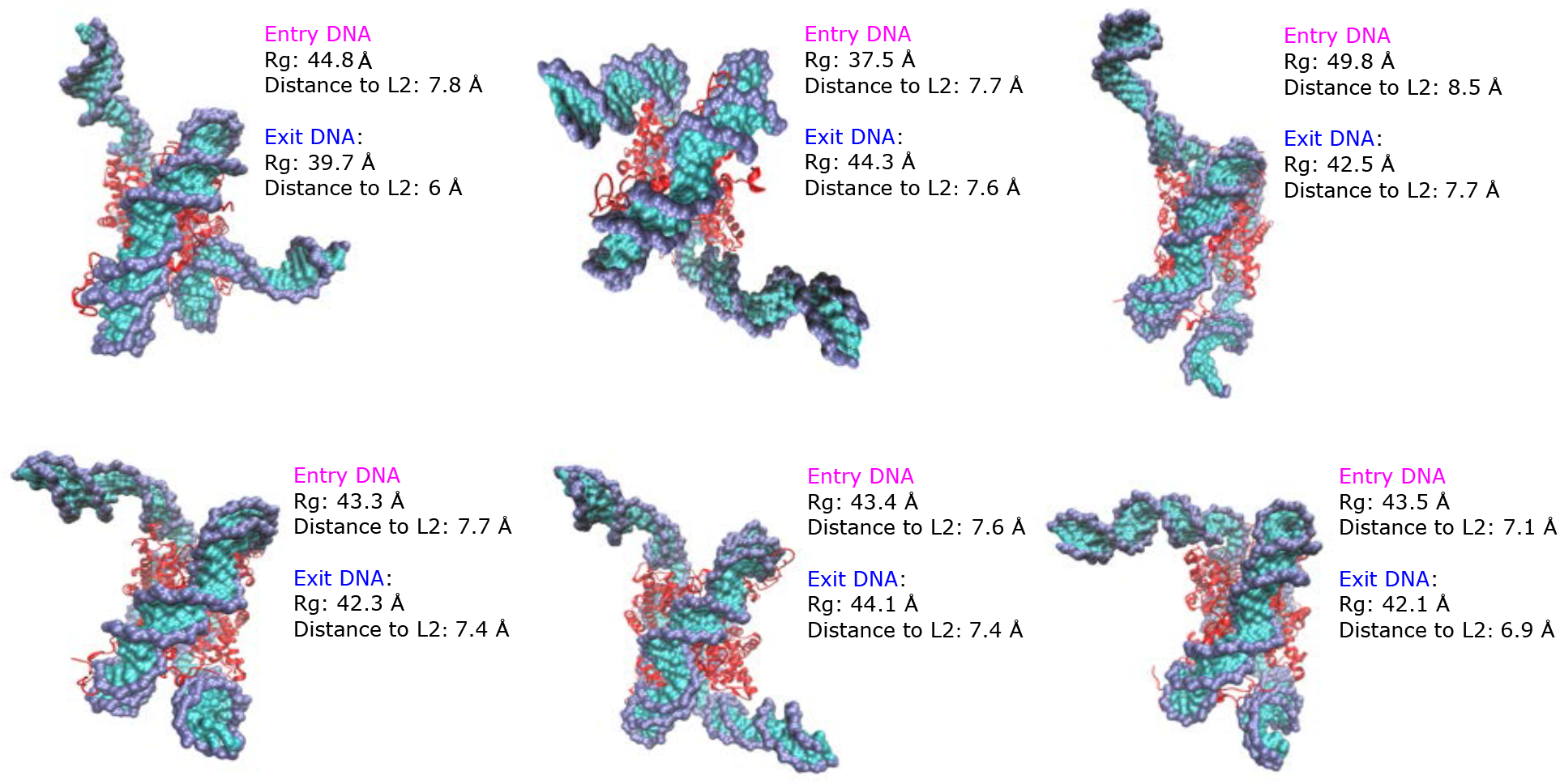
Structural clusters of H2A nucleosomes extracted from ABMD simulations. Structures represent close and semi-close states with raddi of gyration between 37.5 Å to 49.8 Å for entry DNA and from 39.7 Å to 44.3 Å for exit DNA. Distance to L2 shows the distance of the L2-loop residues to their adjacent DNA base pairs. Compared to H2A.B NCPs, the DNA L2-loop distances are lower which is in agreement with lower radii of gyration. See Figs. S7 & S8 for side views of these clusters.

**Figure 8:**
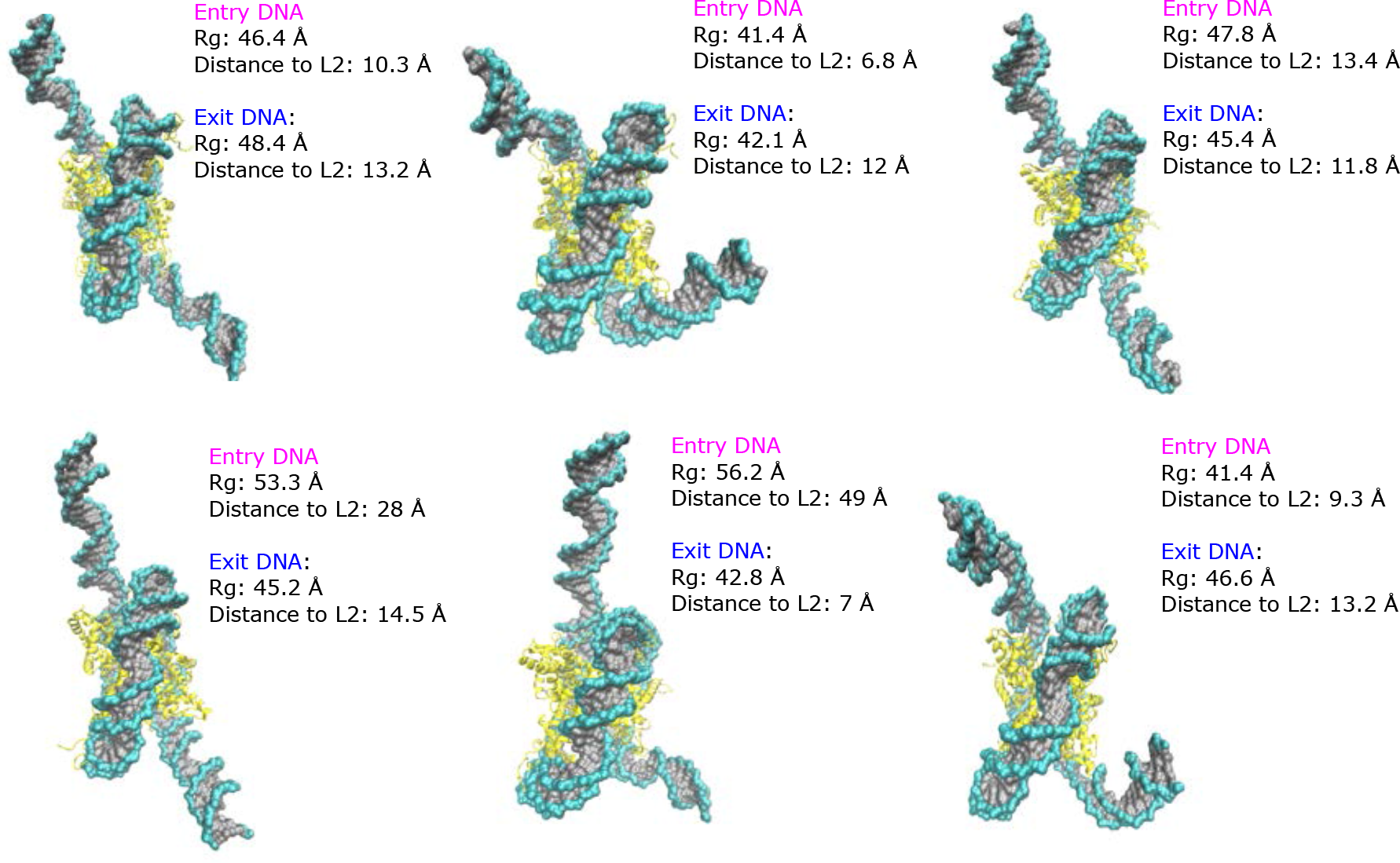
Structural clusters of H2A.B nucleosomes extracted from ABMD simulations. Structures represent semi-open and open states with raddi of gyration between 41.4 Å to 56.2 Å for entry DNA and from 42.1 Å to 48.4 Å for exit DNA. Distance to L2 shows the distance of the L2-loop residues to their adjacent DNA base pairs. Increasing radii of gyration are associated with increased DNA L2-loop distances in the open states of H2A.B NCPs. See Figs. S9 & S10 for side views of these clusters.

### H2A.B Reduces Long Range Dynamic Correlations

In previous simulations of canonical and macroH2A nucleosomes, we found that dynamic networks propagate throughout the system using a combination of distinct protein and DNA networks (65). To determine the effects of H2A.B incorporation on these networks, the mutual information of the protein C_*α*_ and DNA C1’ atom dynamics were computed for canonical and H2A.B systems in open and closed states (Fig. S13). For both systems, long range correlations were decreased upon DNA opening from the compact to the open states. This includes protein-protein correlations, indicating that DNA wrapping around the histone core helps to stabilize networks within it. Differences between the canonical and H2A.B systems show that in both the compact and open states, H2A.B significantly reduces long range correlations in both the DNA and protein core. In particular, the difference between the two closed state systems shows that correlations are stronger in the DNA and H3 proteins in canonical nucleosomes relative to H2A.B systems (Fig. 9a). For open states there is a more dramatic decrease in correlations upon H2A.B replacement, particularly for residues in the H3/H4 tetramer (Fig. 9b). Given that canonical and H2A.B nucleosomes favor the compact and open states respectively, we also computed the correlation difference between the compact H2A and open H2A.B systems (Fig. 9c). Here, there was a substantial increase in correlations throughout the system by moving from the open H2A.B nucleosome to the closed canonical state. In general, these results show that H2A.B decreases long range correlations and weaken allosteric networks in the nucleosome through a combination of increased DNA opening and changes to the dimer/tetramer and dimer/DNA interfaces.

**Figure 9:**
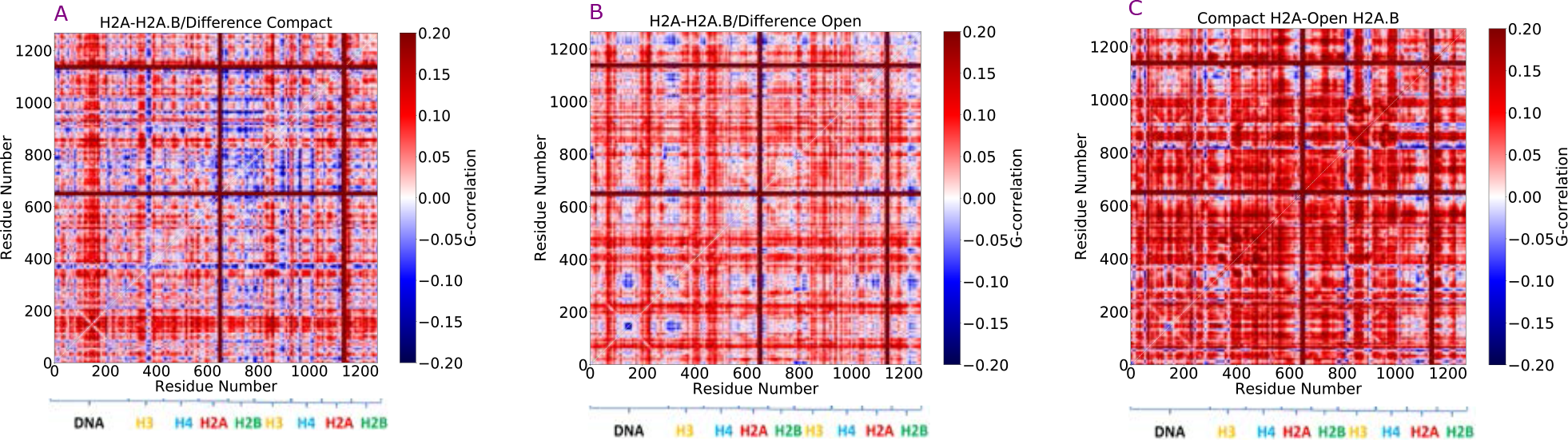
Differences between atomic mutual information calculations of the H2A and H2A.B NCPs in the compact (a) and open states (b). In both cases, there are higher correlated motions in the H2A NCP. (c) The difference between the compact structure of the H2A NCP to the open H2A.B NCP state shows significant stronger correlations in the closed H2A state.

## DISCUSSION

In this work, we used a combination of conventional and enhanced sampling simulations to discern the effects of H2A.B substitution on nucleosome structures and dynamics. Our results show that H2A.B primarily alters intra-nucleosome interactions in two regions: the DNA-dimer and dimer-tetramer interfaces. At the DNA-dimer interface, residues in the H2A.B L2 loops appear to have the most significant influence on DNA stability, as the canonical Lys-Thr-Arg sequence is replaced with Glu-Arg-Asn in H2A.B. This substitution reduces the L2 loop’s net charge from +2 to 0 and its ability to form stabilizing hydrogen bonds with the DNA. As we observed in enhanced sampling simulations, this point of contact was routinely broken in H2A.B nucleosomes, resulting in sampling of more elongated states with DNA opening by up to 20 basepairs on each side. These open state are stable on the hundreds of nanonseconds timescales, as exemplified by our conventional simulations of the dominant structural clusters. This central role of the L2 loops was highlighted in a recent complementary molecular dynamics study of H2A.B nucleosomes, in which Peng *et al*. observed a similar role for these residues (80). This change to the L2 loops appears to be unique to the H2A.B variant, as macroH2A, H2A.X, and H2A.Z all have two basic and no acidic residues in this sequence (11). Therefore, mutations targeting the basic residues in the L2 loops are likely to have significant effects on the free energy difference between open and closed nucleosome states.

The effects of H2A.B modifications at the dimer/tetramer interface on nucleosome structures are less clear. Although an energetic analysis showed a significantly weaker interface at the docking domain, which is shorter and less basic in H2A.B, all conventional and enhanced sampling simulations performed here showed the nucleosome core remaining intact with relatively minor structural rearrangements. It has been proposed that dimers can exhibit a butterfly opening motion which would occur on timescales that are far beyond what are sampled here (81). Furthermore, fluorescence recovery after photobleaching (FRAP) experiments have shown that H2A.B increases the rate of dimer loss and exchange *in vivo* (82). The weaker dimer/tetramer interactions we noted may results in enhancement of these butterfly motions and are likely one of the driving forces of this increased dimer exchange, which may further enhance transcription and nucleosome disassembly processes which utilize subnucleosomal particles (83). However, at equilibrium, our simulations, as well as previously performed SAXS experiments(28), suggest that dominant conformation of H2A.B systems *in vitro* are intact nucleosome cores with DNA existing in a dynamic equilibrium of open states. This weaker interface may also make H2A.B systems more susceptible to disassembly by post-translational modifications or changes in solution conditions (67, 84, 85). Finally, while all simulations presented here used the same DNA sequence, the fact that H2A.B nucleosomes are less capable of wrapping DNA than their canonical counterparts suggests that DNA sequences may have disparate effects in these two systems. In particular, given that H2A.B systems favor open DNA states, sequences which are less rigid and more likely to adopt highly bent nucleosome states, such as Widom 601, may preferentially bind to canonical containing nucleosomes.

The decrease of long range dynamic correlations throughout the nucleosome observed here, which is a result of both DNA unwrapping and H2A.B substitution altering protein/protein and protein/DNA contacts, may play an additional role in epigenetic regulation. Indeed, allosetric networks within the nucleosome have been observed with solid state NMR experiments (86), and it has been proposed that epigenetic regulators may take advantage of these networks to influence genetic activity (87). Furthermore, it has been shown that allosteric transmission can be exploited to design drugs that bind at sites on the nucleosome surface (88), and that these networks can extend between nucleosomes to alter chromatin fiber dynamics (89). Therefore, further analyses is warranted on the influence of H2A.B on chromatin dynamics and how these dynamics may alter genetic accessibility far from the H2A.B substitution site.

## AUTHOR CONTRIBUTIONS

H.K. performed the simulations. H.K. and J.W. designed the experiments, analyzed data, and wrote the manuscript.

## ACKNOWLEDGMENTS

The authors thank Dr. Samuel Bowerman, Mr. Joseph Clayton, and Mr. Dustin woods for valuable discussions concerning this work. This research was supported by the National Institute of General Medical Sciences of the National Institutes of Health (Grants 1R15GM114758 & 1R35GM119647). The content is solely the responsibility of the authors and does not necessarily represent the official views of the National Institutes of Health.

## Notes

### Competing Interest Statement

The authors have declared no competing interest.

