## Supporting Material for "Effects of H2A.B incorporation on nucleosome structures and dynamics"

### Supplemental Material

Havva Kohestani[1] and Jeff Wereszczynski[2]

[1]Department of Biology and the Center for the Molecular Study of Condensed Soft Matter, Illinois Institute of Technology, Chicago, IL 60616
[2]Department of Physics and the Center for the Molecular Study of Condensed Soft Matter, Illinois Institute of Technology, Chicago, IL 60616

### S1 MM/GBSA INPUT FILES

Here we supply an example input files for the *mmpbsa.py* program, which we have used for estimating interaction energies:

```
Per-residue MM/GBSA decomposition
&general
  endframe=100, verbose=1, keep_file=2/
&gb
  igb=5, saltcon=0.150,/
&decomp
  idecomp=1, print_res="1-706", csv_format=0, dec_verbose=0/
```

| Simulation Protocol | NCP System | Number of Replica | Time | Total |
|---|---|---|---|---|
| cMD of Compact States | H2A | 2 | 0.55 μs | 1.1 μs |
| | H2A.B | 2 | 0.55 μs | 1.1 μs |
| cMD of Open States | H2A | 12 | 0.55 μs | 6.6 μs |
| | H2A.B | 12 | 0.55 μs | 6.6 μs |
| Enhanced MD | H2A | 3 | 1.6 μs | 4.8 μs |
| | H2A.B | 3 | 1.6 μs | 4.8 μs |

Table S1: Summary of conventional and enhanced sampling simulations performed in this study.

| System | Interaction | SASA $\Delta E$ | Cavity & Dispersion $\Delta E$ | $\Delta\Delta E$ |
|---|---|---|---|---|
| Canonical | Dimer-Tetramer | -23.7±1.7 | -29.1±2.8 | 5.4±3.3 |
| | H2A-Tetramer | -13.7±1.4 | -16.4±2.3 | 2.7±2.7 |
| | Dimer-DNA | -37.3±1.4 | -45.7±1.9 | 8.4±2.4 |
| H2A.B | Dimer-Tetramer | -16.2±1.7 | -20.9±2.8 | 4.7±3.3 |
| | H2A-Tetramer | -7.7±1.5 | -9.3±2.3 | 1.6±2.7 |
| | Dimer-DNA | -34.7±1.3 | -43.6±1.9 | 8.9±2.7 |

Table S2: Comparison of the non-polar interaction energies in H2A and H2A.B nucleosomes calculated based on the solvent exposed surface area (SASA) and Cavity & Dispersion term methods. Overall energy differences between the two methods are small relative to the interaction energies report in Table 1. Default values were used for all calculations.

| MM/GBSA | *igb=5, mbondi=2* | | | |
|---|---|---|---|---|
| System | Interacting Segments | $\Delta E_{vdW}$ | $\Delta E_{elec}$ | $\Delta E_{total}$ |
| HA2 NCP | Dimer-Tetramer | -226.5±6.6 | 108.8±36.6 | -118.5±11.6 |
| | H2A-Tetramer | -127.7±5.8 | 64.4±24.2 | -63.3±15 |
| | Dimer-DNA | -368.9±6.4 | -11.9±3.8 | -380.8±5.1 |
| H2A.B NCP | Dimer-Tetramer | -169.9±6.3 | 88.1±38 | -81.8±22.1 |
| | H2A.B-Tetramer | -77.6±5.4 | 33.1±24.1 | -44.5±14.7 |
| | Dimer-DNA | -357.1±6.2 | -14.9±5.8 | -372±9.3 |
| MM/GBSA | *igb=8, mbondi=3* | | | |
| System | Interacting Segments | $\Delta E_{vdW}$ | $\Delta E_{elec}$ | $\Delta E_{total}$ |
| HA2 NCP | Dimer-Tetramer | -238.2±6.1 | 113.3±4.8 | -124.9±5.1 |
| | H2A-Tetramer | -137.9±4.7 | 55.2±3.6 | -82.7±4.4 |
| | Dimer-DNA | -369.5±7.2 | -27.9±10.6 | -397.4±11.2 |
| H2A.B NCP | Dimer-Tetramer | -173.9±4.6 | 83.9±4.8 | -90±19.1 |
| | H2A.B-Tetramer | -79.8±3.1 | 32.1±3.7 | -47.7±3.4 |
| | Dimer-DNA | -357.7±8.8 | -22.7±9.2 | -380.4±10.6 |

Table S3: Energy differences (kcal/mol) of association between the dimer/tetramer, H2A/tetramer, and dimer/DNA constituents as estimated by an MM/GBSA analysis calculated for *igb*=5 and *igb*=8. A negative value indicates more favorable interactions in the complex relative to in solution. The total interaction energy ($\Delta E_{total}$) is a sum of the van der Waals ($\Delta E_{vdW}$) and electrostatic ($\Delta E_{elec}$) interactions.

| Time Scale | 250 ns | | | |
|---|---|---|---|---|
| System | Interacting Segments | $\Delta E_{vdW}$ | $\Delta E_{elec}$ | $\Delta E_{total}$ |
| HA2 NCP | Dimer-Tetramer | -234.2±10.5 | 112.2±5.5 | -122±9.9 |
| | H2A-Tetramer | -134.5±6.3 | 55±9.5 | -79.5±7.9 |
| | Dimer-DNA | -366.8±6.2 | -21.9±13.9 | -388.7±10.1 |
| H2A.B NCP | Dimer-Tetramer | -172±9.1 | 85.6±6.5 | -86.4±17.5 |
| | H2A.B-Tetramer | -78.8±6.4 | 34.7±5.8 | -44.1±5.5 |
| | Dimer-DNA | -349.7±6.8 | -20.8±7.6 | -370±15.9 |
| Time Scale | 500 ns | | | |
| System | Interacting Segments | $\Delta E_{vdW}$ | $\Delta E_{elec}$ | $\Delta E_{total}$ |
| HA2 NCP | Dimer-Tetramer | -238.2±6.1 | 113.3±4.8 | -124.9±5.1 |
| | H2A-Tetramer | -137.9±4.7 | 55.2±3.6 | -82.7±4.4 |
| | Dimer-DNA | -369.5±7.2 | -27.9±10.6 | -397.4±11.2 |
| H2A.B NCP | Dimer-Tetramer | -173.9±4.6 | 83.9±4.8 | -90±19.1 |
| | H2A.B-Tetramer | -79.8±3.1 | 32.1±3.7 | -47.7±3.4 |
| | Dimer-DNA | -357.7±8.8 | -22.7±9.2 | -380.4±10.6 |

Table S4: Energy differences (kcal/mol) of association between the dimer/tetramer, H2A/tetramer, and dimer/DNA constituents as estimated by an MM/GBSA analysis calculated for 250 ns and 500 ns with *igb=8, mbondi=3* protocol. A negative value indicates more favorable interactions in the complex relative to in solution. The total interaction energy ($\Delta E_{total}$) is a sum of the van der Waals ($\Delta E_{vdW}$) and electrostatic ($\Delta E_{elec}$) interactions.

| Histone-Residue | $\Delta E_{vdW}$ | $\Delta E_{elec}$ | $\Delta E_{total}$ |
|---|---|---|---|
| H3-ARG 8 | -2.1±1.1 | -1.6±3.8 | -3.8±2.4 |
| H3-ARG 134 | -4.1±1.8 | -0.2±3.5 | -4.3±2.7 |
| H4-ARG 92 | -5.2±1.2 | -1.8±1.3 | -6.9±1.3 |
| H4-TYR 98 | -11.3±1.2 | 0.8±0.3 | -10.2±0.8 |
| H2B-LEU 97 | -6.4±0.7 | 0.5±0.6 | -5.9±0.6 |
| H2A-ARG 99 | -3.2±1.3 | -2.1±3.4 | -5.2±2.4 |
| H2A-THR 101 | -5.5±0.6 | 1.2±0.4 | -4.3±0.5 |
| H2A-ALA 103 | -2.9±0.3 | -0.8±0.1 | -3.7±0.2 |
| H2A-LEU 115 | -4.9±0.8 | -1.3±0.5 | -6.3±0.6 |
| H2A-GLU 121 | -2.5±1.5 | -1.2±6.7 | -3.8±4.1 |

Table S5: Residues with the highest energy differences (kcal/mol) of association at the dimer/tetramer interface for the H2A NCP as estimated by an MM/GBSA analysis. A negative value indicates more favorable interactions in the complex relative to in solution. The total interaction energy ($\Delta E_{total}$) is a sum of the van der Waals ($\Delta E_{vdW}$) and electrostatic ($\Delta E_{elec}$) interactions.

| Histone-Residue | $\Delta E_{vdW}$ | $\Delta E_{elec}$ | $\Delta E_{total}$ |
|---|---|---|---|
| H3-GLU 94 | -2.6±0.5 | 3.5±1.7 | 0.9±1.1 |
| H3-GLU 97 | -0.1±0.1 | 0.7±1.1 | 0.5±0.5 |
| H3-ASN 108 | -1.2±0.2 | 1.9±0.2 | 0.7±0.2 |
| H4-LYS 44 | -1.8±0.6 | 4.3±2.6 | 2.5±1.6 |
| H4-ASP 68 | -1.7±0.3 | 5.6±1.3 | 3.8±0.8 |
| H4-ASP 85 | -0.1±0.2 | 1.4±1.7 | 1.2±0.5 |
| H4-LYS 91 | -1.3±0.8 | 1.8±3.5 | 0.5±2.1 |
| H2B-GLU 73 | -0.6±0.9 | 3.5±1.4 | 2.9±1.2 |
| H2A-LYS 118 | -0.6±0.4 | 1.6±7.2 | 0.9±3.8 |
| H2A-LYS 119 | -0.2±0.2 | 0.8±4.2 | 0.6±2.2 |

Table S6: Residues with the lowest energy differences (kcal/mol) of association at the dimer/tetramer interface for the H2A NCP as estimated by an MM/GBSA analysis. A negative value indicates more favorable interactions in the complex relative to in solution. The total interaction energy ($\Delta E_{total}$) is a sum of the van der Waals ($\Delta E_{vdW}$) and electrostatic ($\Delta E_{elec}$) interactions.

| Histone-Residue | $\Delta E_{vdW}$ | $\Delta E_{elec}$ | $\Delta E_{total}$ |
|---|---|---|---|
| H4-ARG 92 | -4.3±0.9 | -1.4±4.3 | -5.7±2.6 |
| H4-LEU 96 | -3.1±0.3 | -0.3±0.8 | -3.4±0.5 |
| H4-TYR 98 | -11.2±1.1 | 0.7±1.3 | -10.5±1.1 |
| H4-GLY 99 | -3.2±0.5 | -0.1±3.1 | -3.3±1.8 |
| H2B-SER 61 | -3.1±0.7 | 0.1±2.3 | -3.1±1.5 |
| H2B-TYR 80 | -5.5±0.7 | 2.3±1.1 | -3.2±0.9 |
| H2B-LEU 97 | -5.7±0.7 | 0.2±1.9 | -5.4±1.3 |
| H2B-THR 100 | -3.3.1±0.6 | 0.1±1.1 | -3.1±0.8 |
| H2A.B-THR 101 | -4.7±0.8 | 0.9±1.1 | -3.7±0.9 |
| H2A.B-SER 103 | -2.3±0.9 | -3.5±1.7 | -5.9±1.3 |

Table S7: Residues with the highest energy differences (kcal/mol) of association at the dimer/tetramer interface for the H2A.B NCP as estimated by an MM/GBSA analysis. A negative value indicates more favorable interactions in the complex relative to in solution. The total interaction energy ($\Delta E_{total}$) is a sum of the van der Waals ($\Delta E_{vdW}$) and electrostatic ($\Delta E_{elec}$) interactions.

| Histone-Residue | $\Delta E_{vdW}$ | $\Delta E_{elec}$ | $\Delta E_{total}$ |
|---|---|---|---|
| H3-GLU 97 | -0.3±0.1 | 1.2±1.3 | 0.8±0.7 |
| H3-GLU 104 | -1.8±0.6 | 3.9±4.2 | 2.1±2.4 |
| H3-ASP 106 | -0.1±0.1 | 0.8±1.8 | 0.7±0.9 |
| H4-ASP 68 | -1.5±0.2 | 3.7±1.1 | 2.2±0.6 |
| H4-ASP 85 | -0.1±0.1 | 1.3±0.9 | 1.1±0.4 |
| H4-GLY 102 | -1.9.1±1.2 | 2.6±8.1 | 0.6±4.6 |
| H2B-LYS 54 | -1.1±0.6 | 1.7±2.4 | 0.6±1.5 |
| H2B-GLU 73 | -0.5±0.8 | 2.5±1.8 | 2.1±1.3 |
| H2B-GLU 90 | -0.3±0.5 | 1.1±1.4 | 0.7±0.9 |
| H2A.B-ASP 110 | -1.9±1.2 | 3.5±17.1 | 1.5±9.1 |

Table S8: Residues with the lowest energy differences (kcal/mol) of association at the dimer/tetramer interface for the H2A.B NCP as estimated by an MM/GBSA analysis. A negative value indicates more favorable interactions in the complex relative to in solution. The total interaction energy ($\Delta E_{total}$) is a sum of the van der Waals ($\Delta E_{vdW}$) and electrostatic ($\Delta E_{elec}$) interactions.

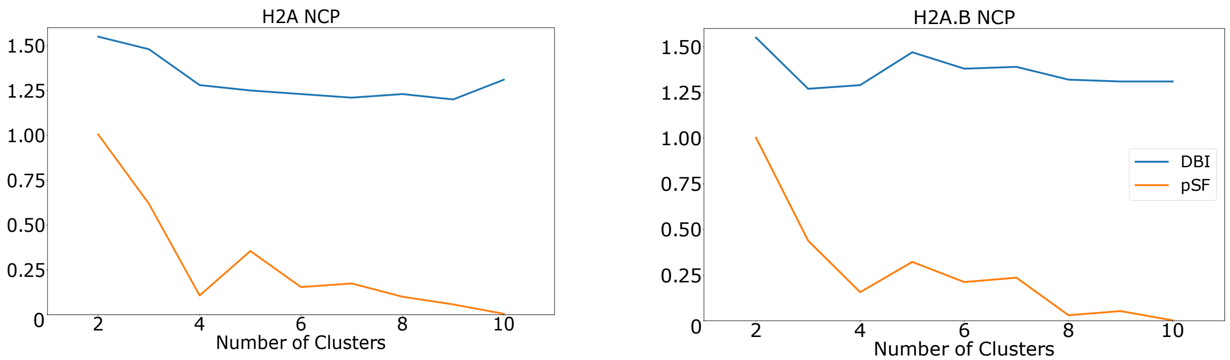


Figure S1: Davis-Bouldin Index (DBI) and seudo-F statistic (pSF) metrics for clustering of ABMD simulations.

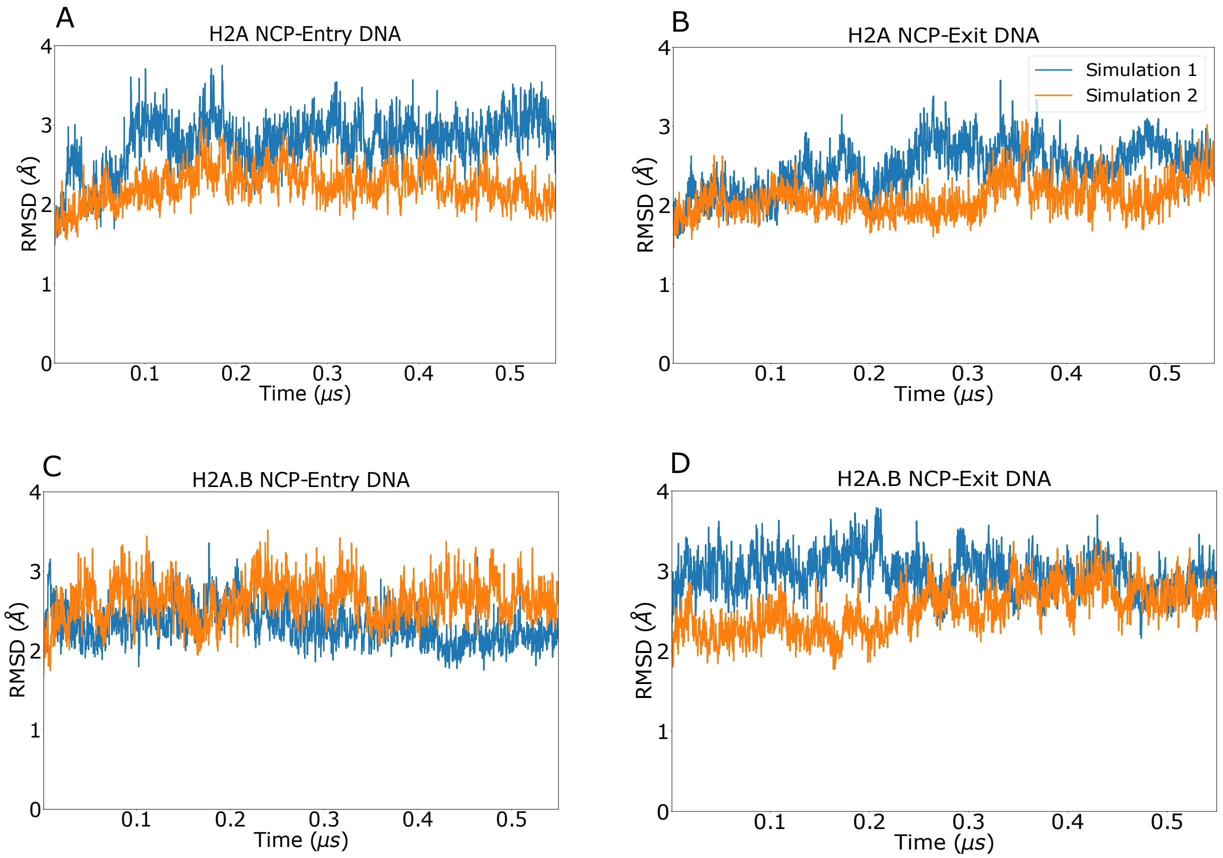


Figure S2: RMSD of entry and exit DNA in H2A and H2A.B nucleosomes in the compact states.

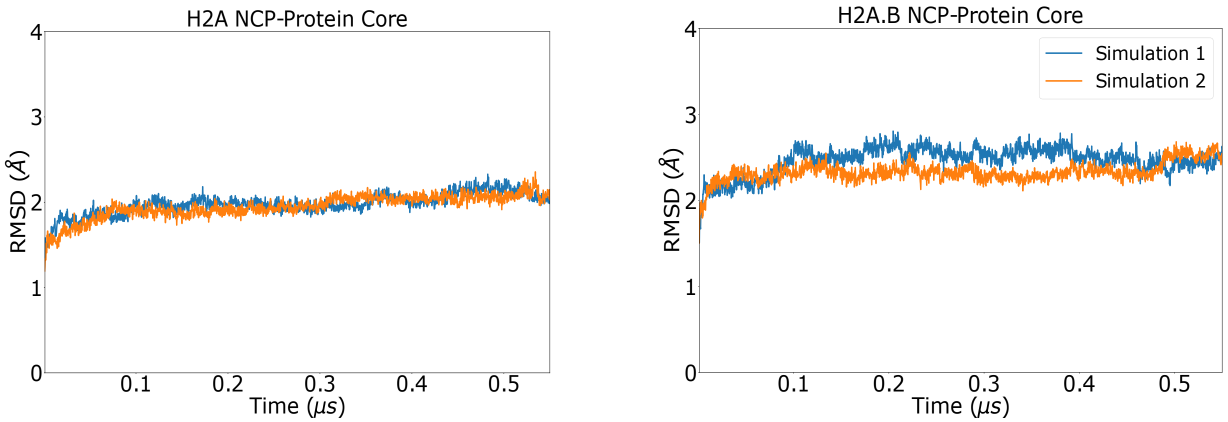


Figure S3: RMSD of the histone core in H2A and H2A.B nucleosomes in the compact states.

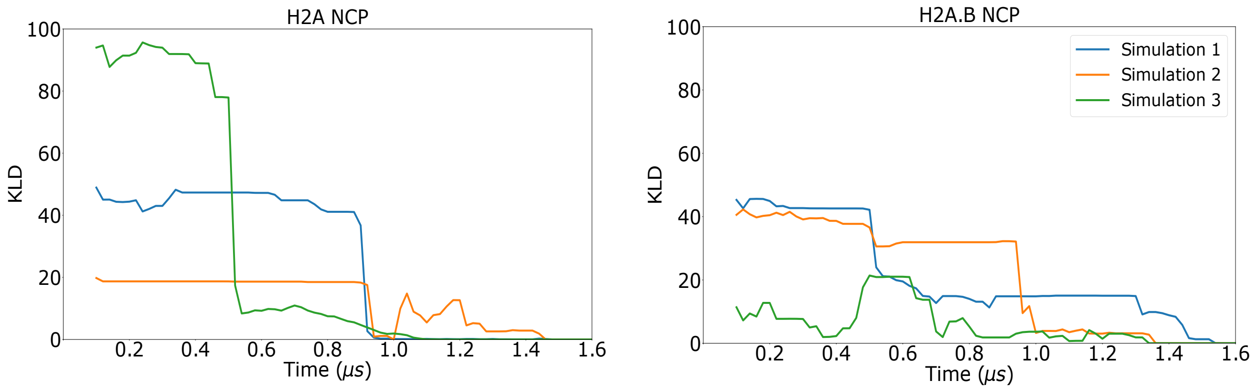


Figure S4: Convergence of the H2A and H2A.B NCPs in ABMD simulation as assessed through a Kullback–Leibler divergence calculation relative to the final sampling probability state (78).

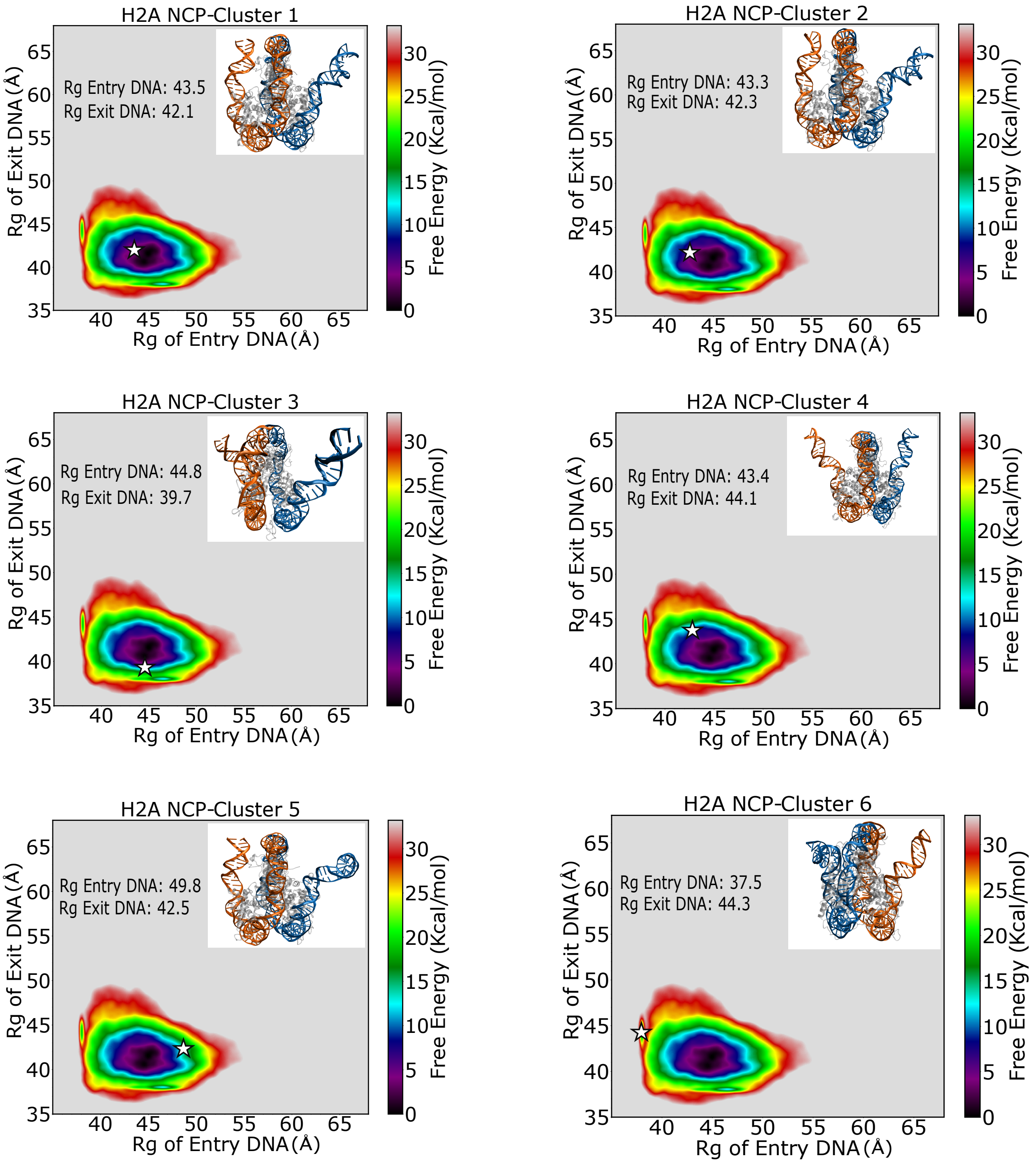


Figure S5: Location of clusters from H2A ABMD simulations on the H2A free energy surface. The clusters are arranged from lowest energy to highest energy. H2A systems sampled primarily compact and semi-open states.

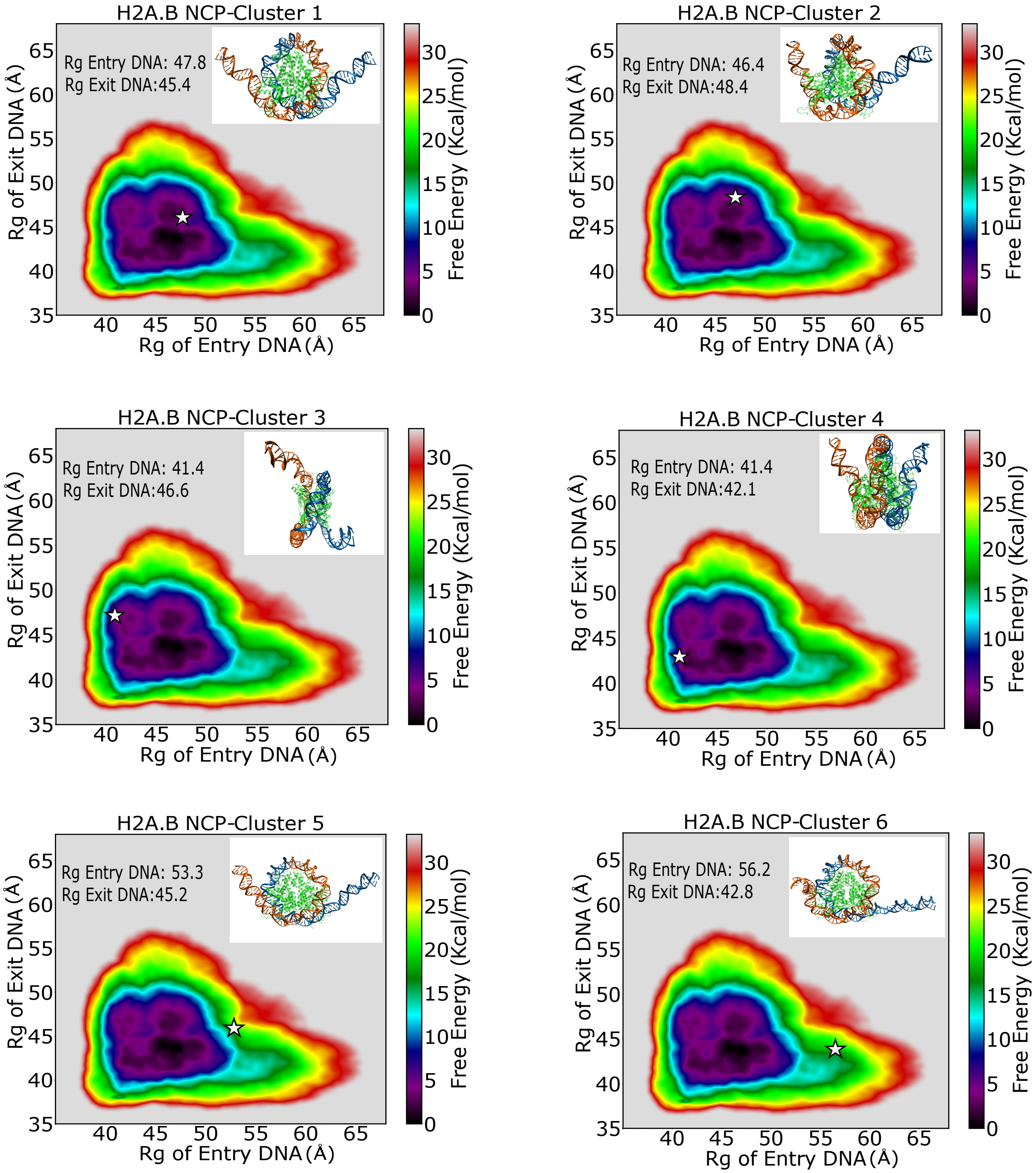


Figure S6: Location of clusters from H2A.B ABMD simulations on the H2A free energy surface. The clusters are arranged from lowest energy to highest energy. H2A.B systems sampled compact through open states.

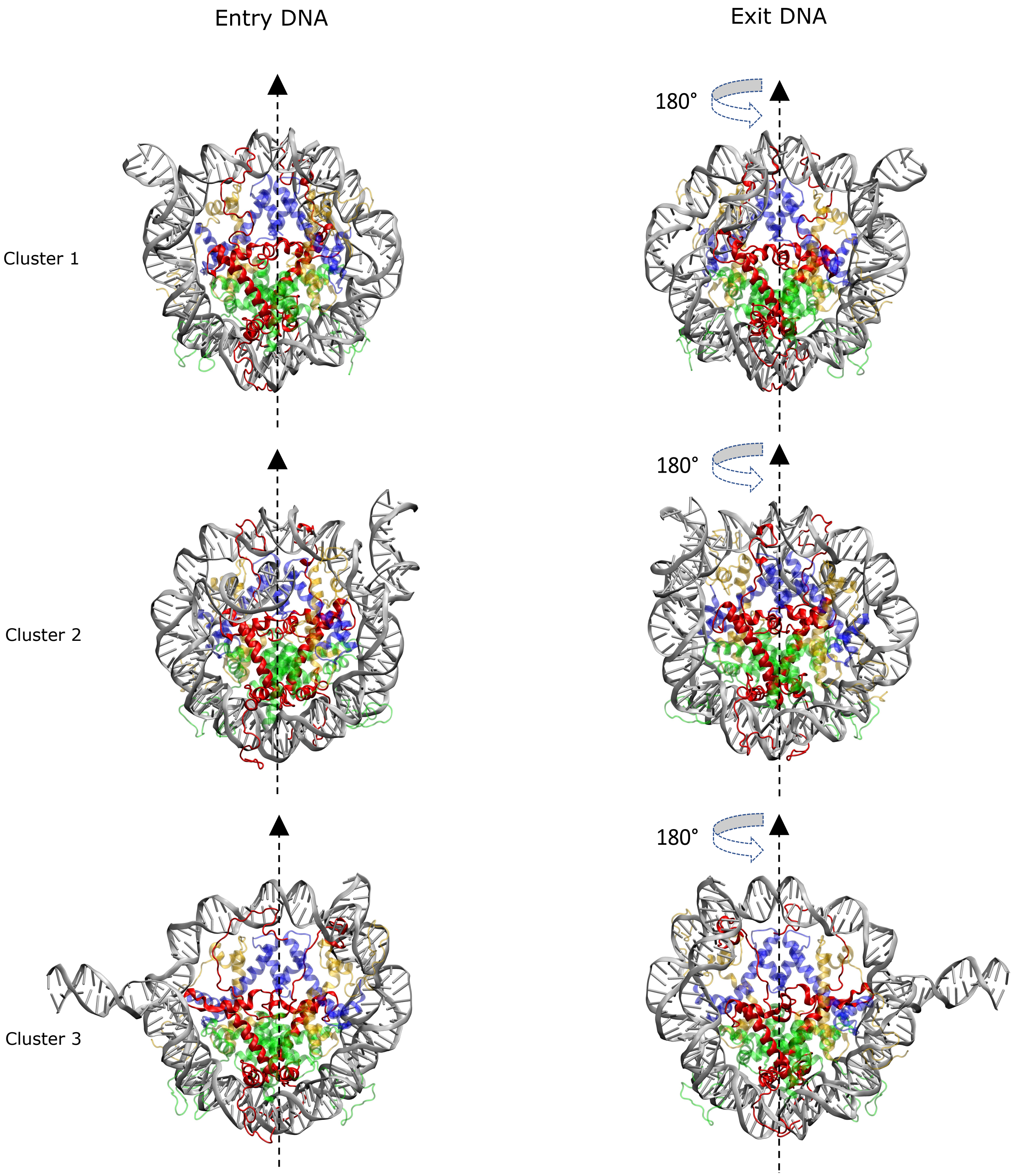


Figure S7: Entry and exit site representations of the H2A nucleosome clusters 1-3 as shown in Figure 7.

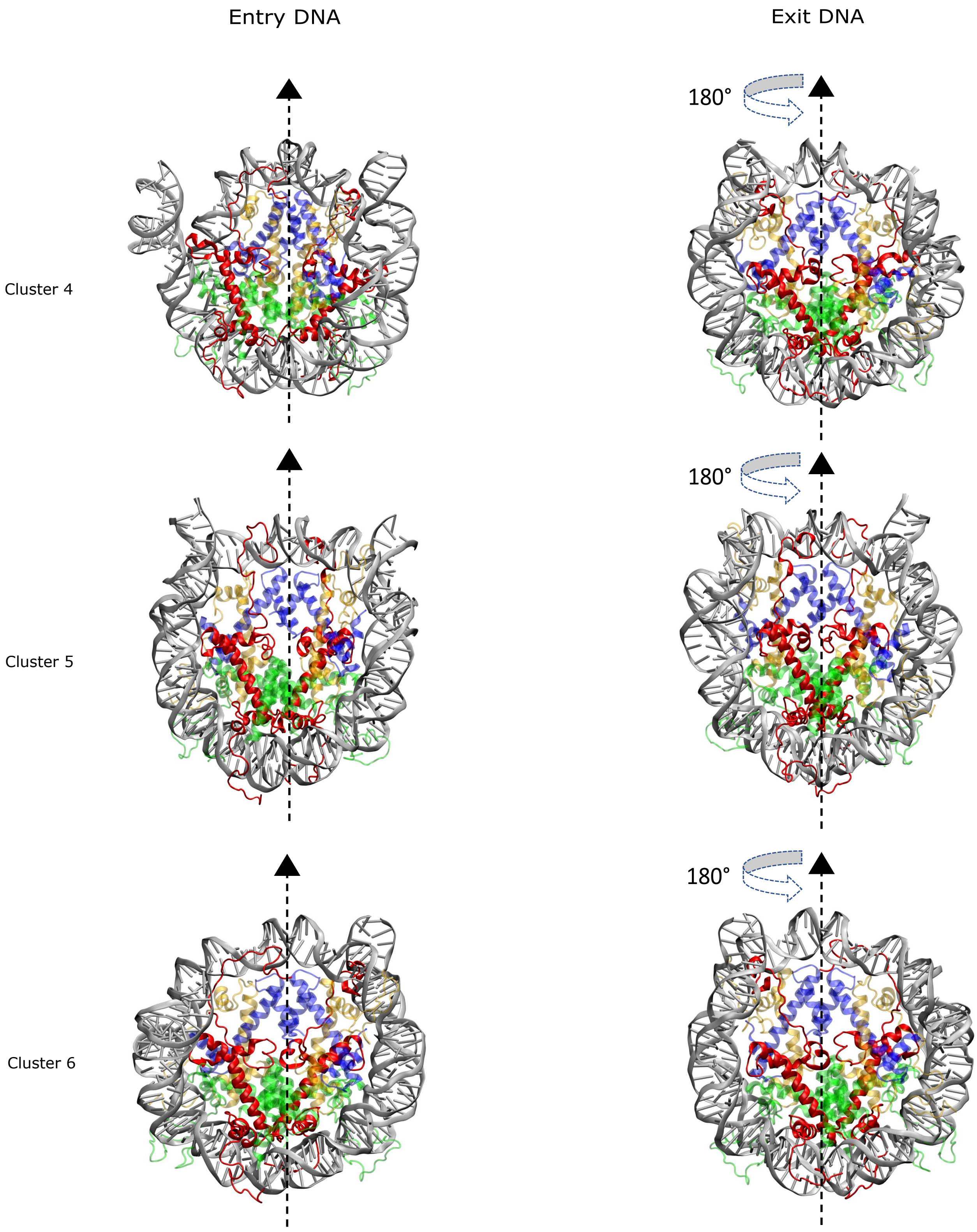


Figure S8: Entry and exit site representations of the H2A nucleosome clusters 4-6 as shown in Figure 7.

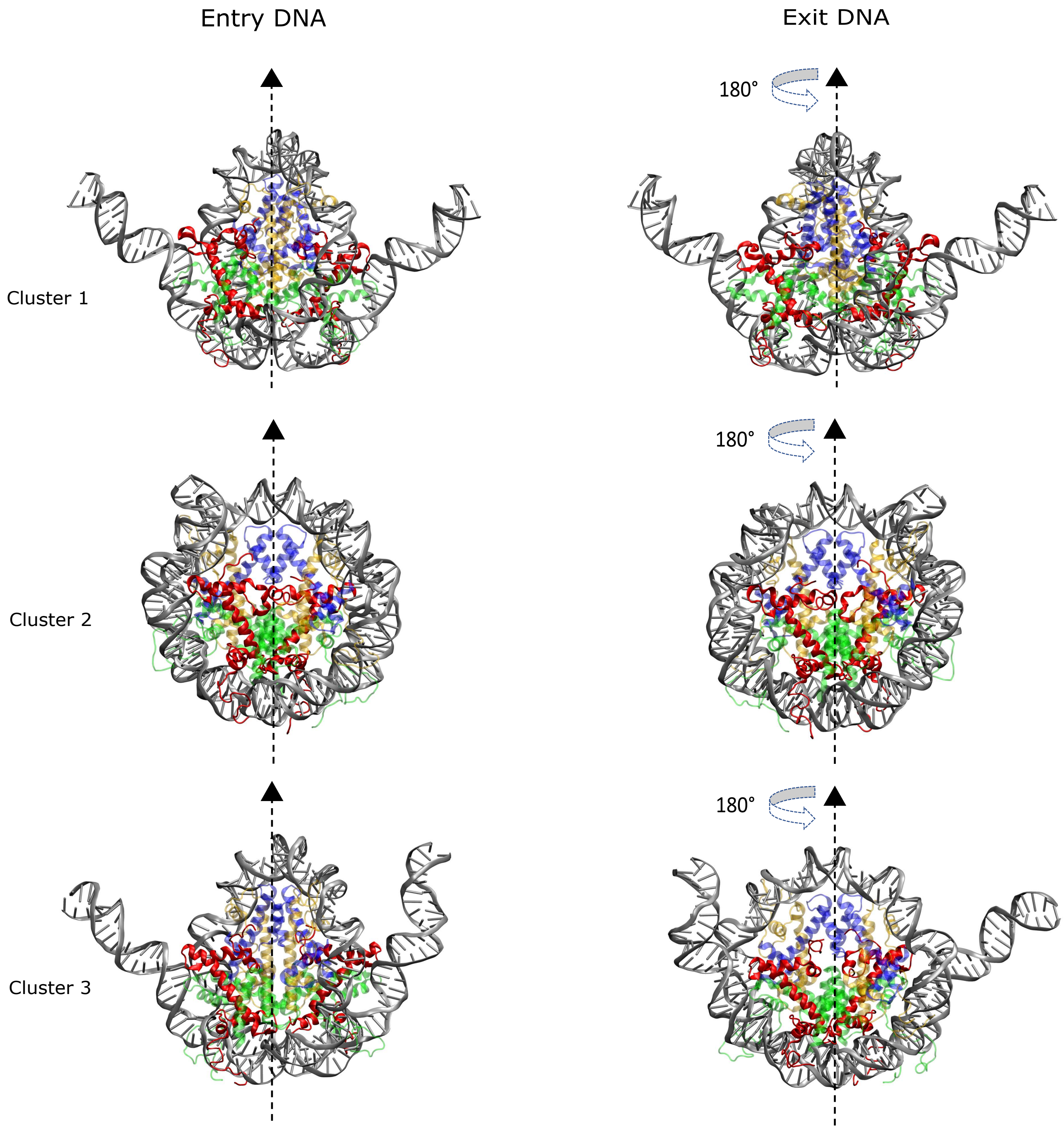


Figure S9: Entry and exit site representations of the H2A.B nucleosome clusters 1-3 as shown in Figure 8.

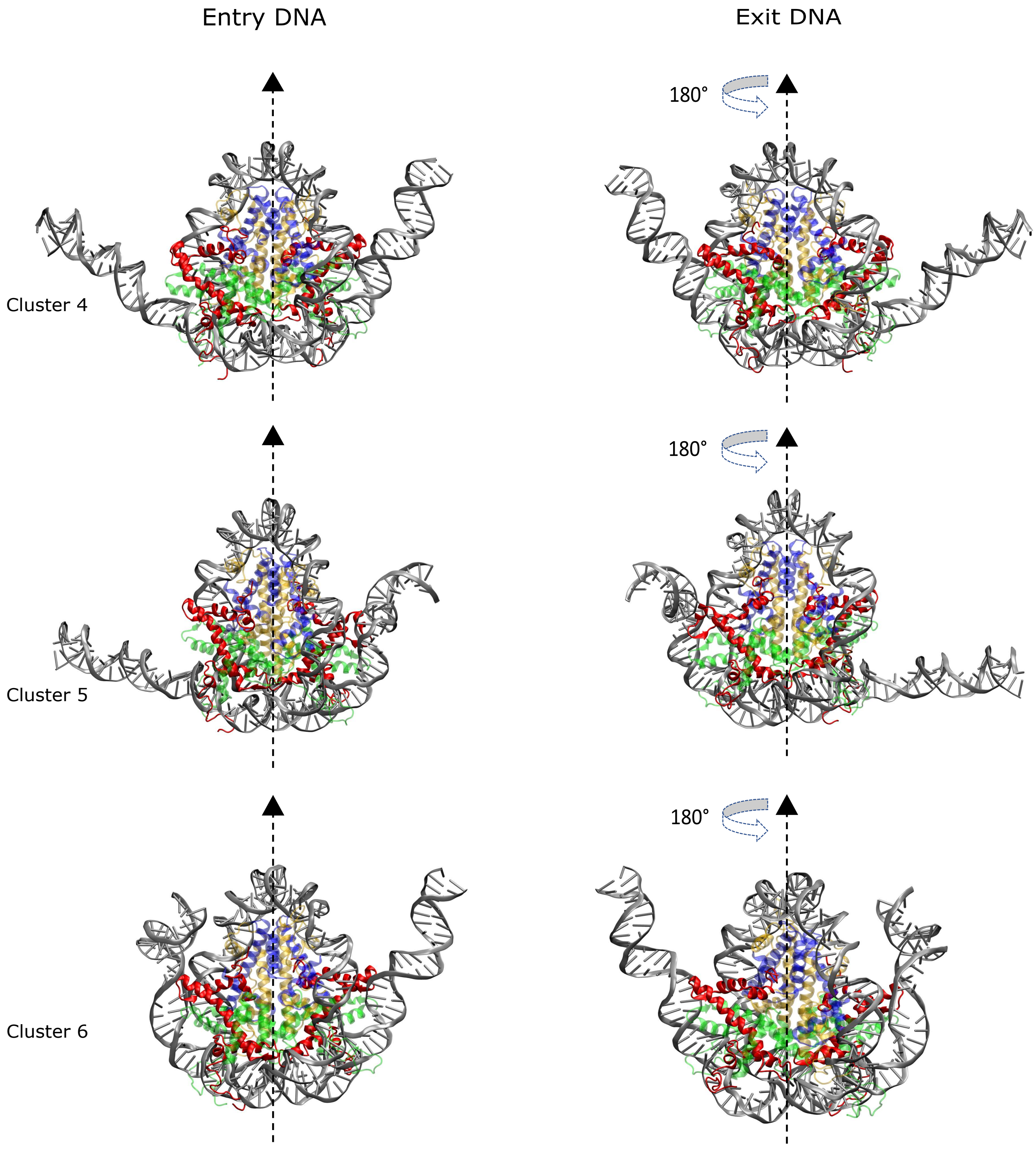


Figure S10: Entry and exit site representations of the H2A.B nucleosome clusters 4-6 as shown in Figure 8.

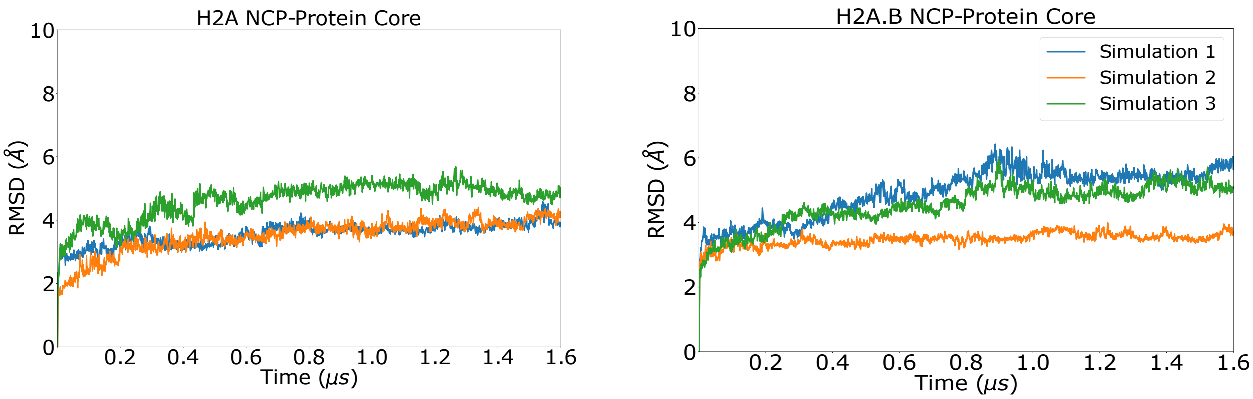


Figure S11: RMSD of histone protein core in H2A and H2A.B nucleosomes during ABMD simulations.

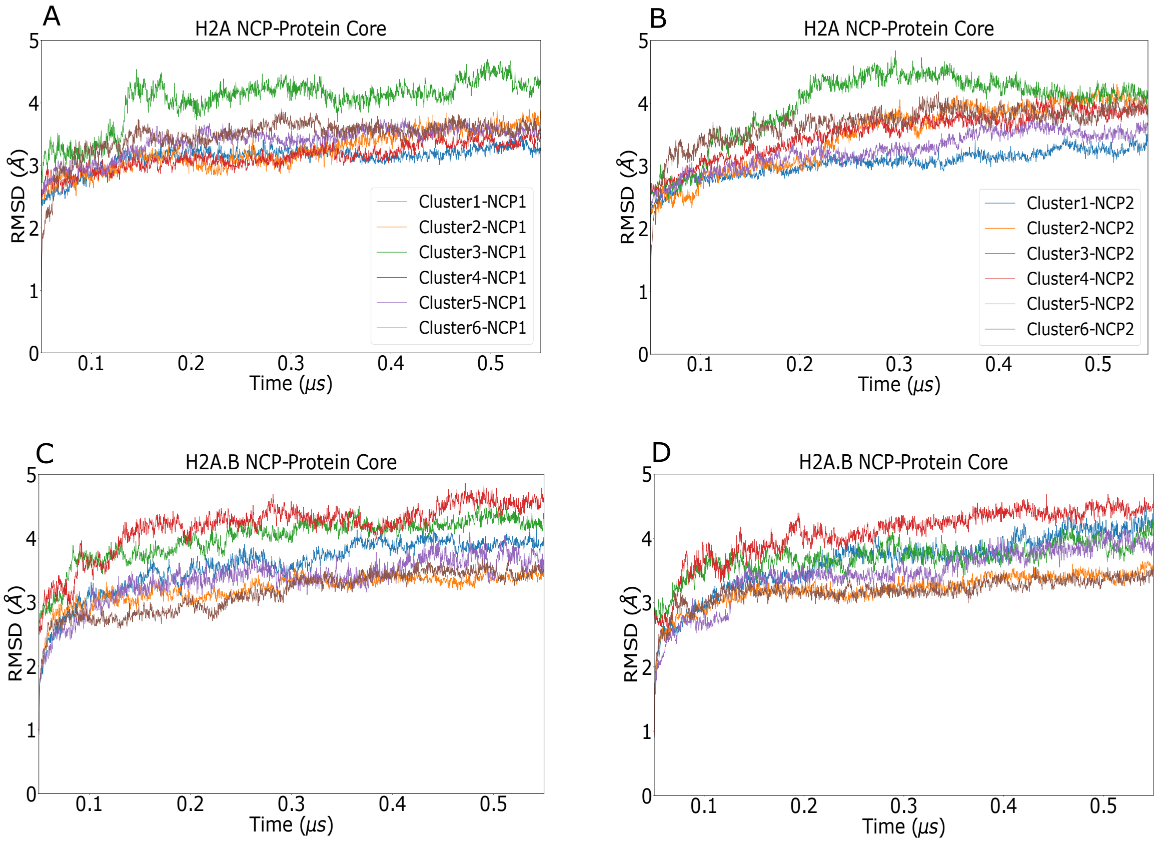


Figure S12: RMSD of histone protein core in H2A and H2A.B nucleosomes during explicit solvent simulations of ABMD structural clusters shown in Figs. 7 & 8. Two copies of each system were simulated, which are denoted as NCP1 and NCP2.

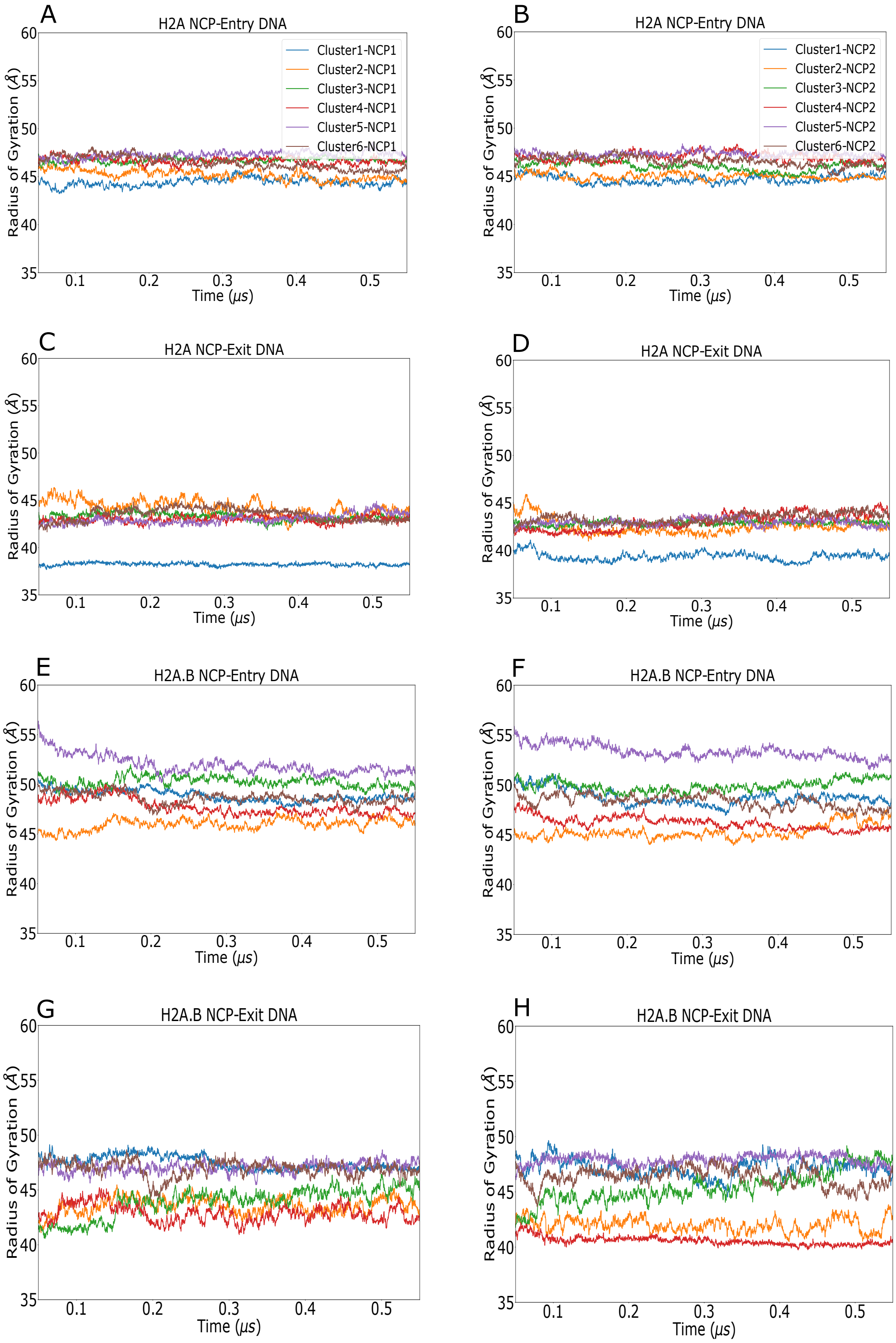


Figure S13: Radius of gyration for entry and exit DNA segments in explicit solvent/open state simulations for H2A (a-d) and H2A.B (e-h) systems from ABMD structural clustering (Figs. 7 & 8). Two copies of each simulation were run (denoted as NCP1 and NCP2). In all simulations the radii of gyration were relatively constant, indicating the sampled open states are stable on the hundreds of nanoseconds timescale.

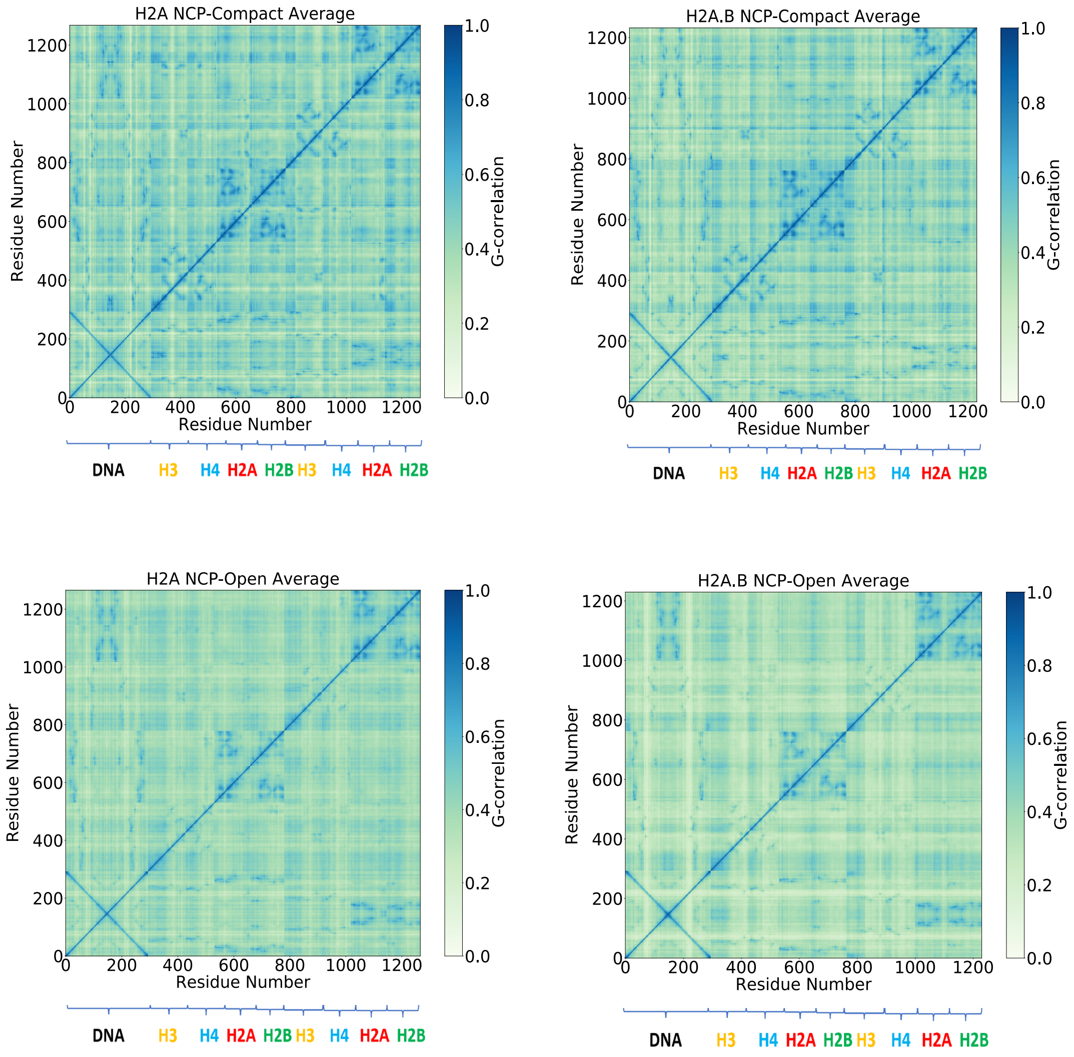


Figure S14: Mutual information calculations of the H2A and H2A.B NCPs averaged for the open and compact states in each system. In both open and compact cases, the H2A NCP has higher correlated motions compared to H2A.B NCP, and the H2A.B NCP in the open state has the weakest overall correlations.
